# Microtubule association induces a Mg-free apo-like ADP pre-release conformation in kinesin-1 that is unaffected by its autoinhibitory tail

**DOI:** 10.1101/2024.11.11.622991

**Authors:** J Atherton, MS Chegkazi, E Peirano, LS Pozzer, T Foran, RA Steiner

**Affiliations:** Randall Centre for Cell and Molecular Biophysics, King’s College London - New Hunt’s House, Guy’s Campus, London SE1 1UL, United Kingdom; Department of Biomedical Sciences, University of Padova, via Ugo Bassi 58/B, Padova, 35131, Italy

## Abstract

Kinesin-1 is a processive dimeric ATP-driven motor that transports vital intracellular cargos along microtubules (MTs). If not engaged in active transport, kinesin-1 limits futile ATP hydrolysis by adopting a compact autoinhibited conformation that involves an interaction between its C-terminal tail and the N-terminal motors domains. Here, using a chimeric kinesin-1 that fuses the N-terminal motor region to the tail and a tail variant unable to interact with the motors, we employed high-resolution cryo-EM in the presence of MTs to investigate elements of the mechanochemical cycle. We describe a missing structure for the proposed two-step allosteric mechanism of ADP release, the ATPase rate limiting step. It shows that MT association induces remodeling of the hydrogen bond network at the nucleotide binding site triggering removal of the Mg^2+^ ion from the Mg^2+^-ADP complex resulting in a strong MT-binding apo-like state before ADP dissociation. We further demonstrate that tail association does not directly affect this mechanism, nor the adoption of the ATP hydrolysis-competent conformation, nor neck linker docking/undocking, even when zippering the two motor domains. Based on this structural evidence, we propose a revised mechanism for tail-dependent kinesin-1 autoinhibition and suggest a possible explanation for its characteristic pausing behavior on MTs.

## Introduction

Kinesin-1 (also known as conventional kinesin) is an essential ATP-driven molecular motor of the kinesin superfamily capable of processive movement as stepping dimers toward the (+)-end of polarized microtubules (MTs)^1^. This enables the transport of a range of cargoes required for cellular function including mitochondria^2,3^, synaptic vesicle and plasma membrane precursors^4^, lysosomes^5,6^, and mRNAs^7^. Several viruses also hijack the MT transport machinery of the host cell to facilitate their replication and spread^8^.

The two motor-bearing heavy chains (Kif5) are typically associated with two light chain (KLC) adaptors that form the tetrameric kinesin-1 complex. KLCs often mediate the interaction with the proteinaceous components of cellular cargoes by recognizing short-linear motifs (SLiMs)^9–11^ and other ordered regions^12^. However, they are not always required as transport of mitochondria and mRNA granules has been shown to be KLC-independent^13,14^. In the case of mitochondria, KLCs even negatively impact on the recruitment of the necessary Milton/Miro adaptor complex that binds directly to Kif5^13^.

Kif5 contains a N-terminal motor domain (also known as the head) that generates force and processive stepping via Mg^2+^-dependent ATP hydrolysis and exchange, multiple coiled-coil (CC) regions with breaks (stalk) that mediate dimerization, and an unstructured C-terminus (tail) involved in autoinhibition (Fig. 1a). ATPase activity is strongly activated by MT binding that induces conformational changes in the motor stimulating both ATP hydrolysis and ADP release steps^15–20^. One of the key structural elements undergoing transitions that are coupled to the nucleotide state is the neck linker, that connects the motor domain to CC1 (Fig. 1a). This element, which is critical for force generation under physiological conditions^21^, goes from being docked along the core motor domain in the ATP-bound state to being flexible in both the ADP-bound and nucleotide-free states^15,20,22–28^. For kinesin-1, MT binding speeds up the rate-limiting step of ADP release more than ∼104 fold^16^. The structural mechanisms governing MT-stimulated ADP release have been disputed for decades and are still unclear^19,23,26,29,30^.

**Figure 1.**
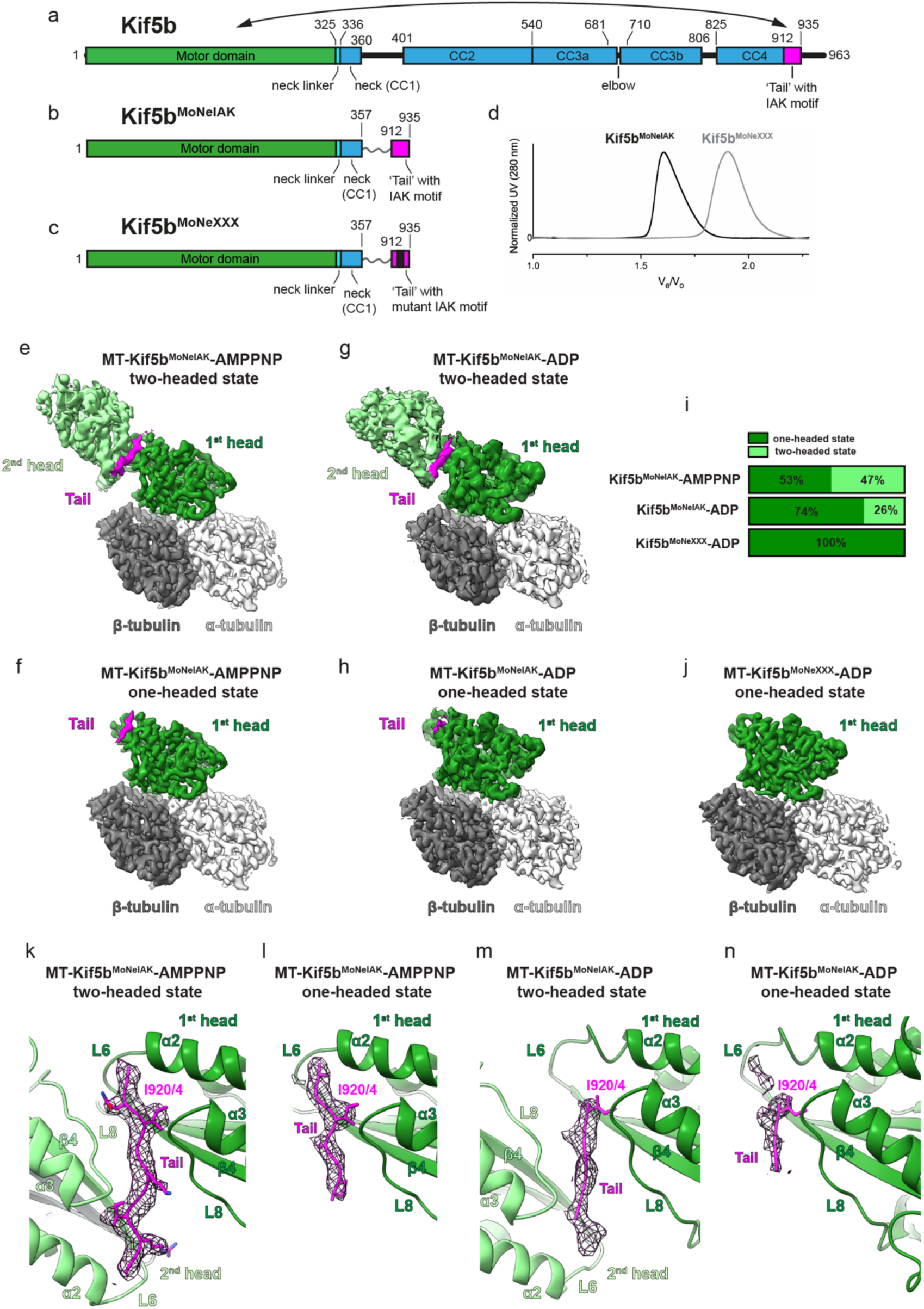
The Kif5 tail IAK motif interacts with MT-bound one-headed and two-headed motors and is required for motor zippering. **a**,**b**,**c.** Schematics showing the domain organization of (**a**) full length Kif5b and of (**b**) Kif5b^MoNeIAK^ and (**c**) Kif5b^MoNeXXX^ chimeras with the mutated IAK motif highlighted in black. Residue numbers are shown for full length Kif5b and this numbering is kept for the chimeras. Kif5b^MoNeIAK^ includes the N-terminal motor domain (head), neck linker and first coiled-coil (neck and CC1) fused by a flexible linker to amino acids 912-935 of the tail domain, containing the IAK motif. **d.** Normalized size-exclusion chromatography (SEC) traces for Kif5b^MoNeIAK^ and Kif5b^MoNeXXX^ (V_e_ = elution volume, V_0_ = exclusion volume). Kif5b^MoNeIAK^ and Kif5b^MoNeXXX^ elute as dimers and monomers, respectively. **e**,**f.** Unsharpened cryo-EM reconstructions of the (**e**) two-headed and (**f**) one-headed states of the MT’s αβ-tubulin heterodimer-Kif5b^MoNeIAK^ asymmetric unit in the presence of AMPPNP. **g**,**h.** Unsharpened cryo-EM reconstructions of the (**g**) two-headed and (**h**) one-headed states of the MT’s αβ-tubulin heterodimer-Kif5b^MoNeIAK^ asymmetric unit in the presence of ADP. **i**. Schematics highlighting the percentage occupancy of one and two-headed states in Kif5b^MoNeIAK^ and Kif5b^MoNeXXX^ datasets. **j.** Overall structure of the MT’s αβ-tubulin heterodimer-Kif5b^MoNeIXXX^ asymmetric unit for which only a one-headed state was observed. **k**,**l.** Cryo-EM density for the Kif5b^MoNeIAK^ tail (mesh) in the MT-Kif5b^MoNeIAK^-AMPPNP (**k**) two-headed and (**l**) one-headed states. **m**,**n**. Cryo-EM density for the Kif5b^MoNeIAK^ tail (mesh) in the MT-Kif5b^MoNeIAK^-ADP (**m**) two-headed and (**n**) one-headed states. In (**k**-**n**) the position of key isoleucine I920/4 and side chains for well-resolved tail residues are shown. L6 and L8 indicate loop 6 and loop 8, respectively.

Another key aspect of kinesin-1’s mechanochemical cycle is its autoinhibition. To limit wasteful ATP hydrolysis and futile runs, cargo-less Kif5 adopts a compacted autoinhibited conformation that involves an interaction between the motor domain and the unstructured C-terminal tail centred on its IAK (isoleucine-alanine-lysine) SLiM^31^ (hereafter just IAK motif, Fig. 1a). Binding of a single tail to the kinesin-1 dimer is sufficient for inhibition^32^ preventing MT-dependent stimulation of ATPase activity, particularly, the acceleration of ADP release^33^. A cryo-EM reconstruction of MT-bound monomeric heads chemically cross-linked to the IAK motif indicated that this inhibitory region engages with the nucleotide-binding switch I motif of loop 9 (see Supplementary Fig. 1a for secondary structure elements of the kinesin-1 motor), with this interaction suggested to directly inhibit ADP release^34^. However, a subsequent crystallographic structure of the autoinhibited ADP-bound *Drosophila* kinesin-1 motor dimer showed the IAK motif zipping the heads by binding away from the nucleotide binding site^35^. This work proposed a ‘double-lockdown’ model of autoinhibition, whereby zipping the heads locks them in a conformationally restricted orientation that, in turn, prevents, undocking of the neck linker from the motor that has been suggested to facilitate ADP release^20,24^.

Recent studies have shown that critical to the achievement of the compact autoinhibited conformation is a short break (elbow) in CC3 of Kif5 that allows the molecule to fold on itself such that the unstructured C-terminal tail is in proximity of the motors^36–38^ (Fig. 1a). These studies suggest that portions of the stalk would reduce MT binding in the folded conformation by sterically interfering with the MT-binding surface of at least one of the motor domains. This is supported by the observation that compared to the motor domain in isolation, MT affinity/landing rate of full-length Kif5 is reduced, with or without KLCs, unless autoinhibition is relieved^33,37,39,40^. However, full-length Kif5 still displays a basal level of landing events and, when MT-associated, reduced motility and longer association/run durations compared to truncated tailless motors^37,40^. The ‘stalling’ behavior characteristic of full-length kinesin-1 motility has been shown to depend on the tail/IAK motif^37,39,40^. Furthermore, addition of a tail peptide to dimeric tailless constructs decreases MT-stimulated ATPase and ADP release rates and motility, stalling kinesin in a strongly-MT-associated state^41–44^. These observations suggest that at least some aspects of kinesin-1’s mechanochemistry are independent of purely steric effects and depend on the direct interaction of the tail with the motor.

Using engineered stalkless chimeric constructs of human Kif5, we present here MT-bound high-resolution cryo-EM structures that explain key aspects of kinesin-1’s mechanochemical cycle and autoinhibition. We have visualized an apo-like transition state of MT-bound kinesin-1 with ADP but lacking Mg^2+^. Our structural analysis shows the MT-induced conformational changes that remove the Mg^2+^ ion leading, in turn, to ADP destabilization and, ultimately, its release. Moreover, we show that whilst the autoinhibitory tail can crosslink the motors in a zipped dimer this arrangement is not rigid and, importantly, it does not prevent undocking of the neck linker. Based on our structures, we propose a revised mechanism for kinesin-1 autoinhibition and suggest an explanation for its distinctive stalling behavior.

## Results

### ‘Bonsai’ kinesin-1 chimeras for cryo-EM studies

It is currently unclear if the association of the autoinhibitory tail affects nucleotide-dependent structural transitions of MT-bound kinesin-1. To explore this, we designed a stalk-less ‘bonsai’ kinesin-1 chimera, dubbed Kif5b^MoNeIAK^ (<u>Mo</u>tor-<u>Ne</u>ck-<u>IAK</u> motif), which fuses the N-terminus of human Kif5b (motor domain up to the end of CC1) to its C-terminal IAK motif via a flexible linker (Fig. 1a,b and Supplementary Fig. 1a). The linker is sufficiently long to allow the free interaction of the tail with the motor. Additionally, we generated an IAK-variant chimera in which we replaced residues 919-QIAKPIR-925 with an unrelated amino acid sequence (TGSTSGT) that was predicted to abrogate motor-tail interactions (Fig. 1c and Supplementary Fig. 1b). We refer to this variant as Kif5b^MoNeXXX^. Size-exclusion chromatography (SEC) runs are consistent with Kif5b^MoNeIAK^ and Kif5b^MoNeXXX^ dimers and monomers, respectively (Fig. 1d). This supports the notion that the IAK motif is involved in crosslinking the motors in the autoinhibited state^35^.

### Tail-bound motors can interact with MTs both as monomers and as zipped dimers

We next sought to assess the structural impact of the tail on Kif5b in ADP and ATP nucleotide states. The ATP analogue adenylyl-imidodiphosphate (AMPPNP) or the ADP+Pi analogue ADP-AlF_4_ can be used to stabilize indistinguishable ATP-like conformational states of the Kif5b motor domain on MTs or tubulin^15,23,26^. Therefore, a cryo-EM dataset was collected of taxol-stabilized MTs decorated with Kif5b^MoNeIAK^ in the presence of AMPPNP for comparison with the high-resolution X-ray crystallographic model of ADP-AlF_4_-bound Kif5b in complex with a tubulin dimer and a DARPin (PDB code 4hna)^15^. Moreover, considering that a high-resolution model of the MT-bound Kif5b motor domain in the presence of ADP is not available, we also collected datasets of taxol-stabilized MTs decorated with Kif5b^MoNeIAK^ and Kif5b^MoNeXXX^ in the presence of ADP. Data collection statistics for all datasets are reported in (Supplementary Table 1).

MT segments of thirteen protofilaments were processed with a modified version of the MiRP MT processing pipeline^45^ (see also the Methods section), followed by symmetry expansion, focused extraction, focused 3D classification and focused 3D refinement on the asymmetric unit. This allowed conformational sorting of individual motor-(αβ)tubulin complexes (Fig. 1e-i, Supplementary Fig. 2). For Kif5b^MoNeIAK^ we recognized two distinct populations of MT-bound motors. Irrespective of the nucleotide state, they were present either in a two-headed state with motors zipped by the autoinhibitory tail (Fig. 1e,g) or in a one-headed state with some tail density still visible associated with the MT-bound motor (Fig. 1f,h). The fraction in the two-headed state was more abundant with AMPPNP (47% of the total) compared to ADP (26%) (Fig. 1i). For ADP-Kif5b^MoNeXXX^ we only observed one-headed motors and no density for the tail (Fig. 1i,j). Whilst one-headed Kif5b^MoNeXXX^ is fully consistent with the monomeric population seen in SEC (Fig. 1d), in the case of Kif5b^MoNeIAK^ its one-headed fraction can be rationalized either because of monomerization due to dilution required for cryo-EM grid preparation or because of flexibility in regions of the neck linker and therefore heterogeneity in the position of the CC and distal head (2^nd^ head) compared to the MT-bound one (1^st^ head) making the later invisible due to the averaging process in cryo-EM reconstruction. In all reconstructions, quality and resolution varied within the asymmetric unit (Supplementary Figs. 3-6), with the MT-motor domain interface being resolved best at ∼2.8-3 Å resolution and decreasing to around ∼3.4–4 Å at the motor tip and tail density. In the two-headed states, the resolution dropped rapidly from ∼3.5 Å to ∼8 Å for the distal motor head moving away from the dimer interface/tail. This decrease in resolution is consistent with some variability in position or conformation for the distal head.

Overall, the arrangement of the MT-bound two-headed complex observed here is consistent with the X-ray crystallographic structure of the autoinhibited MT-free *Drosophila melanogaster* kinesin-1 motors (PDB code 2y65)^35^. However, structural superposition of our cryo-EM structures with the crystallographic one using one of the motors as a frame of reference (the MT-bound one for the cryo-EM structures) reveals variability in the relative positioning of the distal head. This is quantified by ∼14° and ∼21° rotations, in opposite directions and around independent axes, for the AMPPNP and ADP and states, respectively (Supplementary Fig. 7). This suggests that, differently from the symmetric crystallographic complex, some variability in the relative orientation of the zippered heads is compatible the presence of the tail.

In Kif5b^MoNeIAK^, density for the tail is visible in all states — though with varying levels of order — in a cleft between loop 8 and a region leading from the C-terminal tip of helix α2 though loop 6 to the N-terminal end of β4 strand of the core motor domain β-sheet (Fig. 1k-n). This location matches that seen in the MT-free crystallographic structure of autoinhibited *Drosophila* kinesin-1^35^. Although our construct contains a 24 aa-long tail region (Fig. 1b), a maximum of nine amino acids (AQIAKPIRP) could be modelled in the two-headed AMPPNP-Kif5b^MoNeIAK^ structure (Fig. 1k). Like in the X-ray structure, we could not define a clear directionality for the pseudo-palindromic IAK motif thus the peptides were modelled in both polarities (Supplementary Fig. 8a – for clarity, only one peptide is shown in Fig. 1k-n). Sitting at the center of the pseudo-palindrome, K922 has the same location in the complex regardless of polarity with either I920 or I924 embedded in a hydrophobic pocket lined by I130 and F128 of strand β4, I119, Y120 and F116 of helix α2 and C174 of loop 8 of the motor domain (Supplementary Fig. 8a,b). Tail binding is not restricted to the two-headed states that offers pseudo-symmetric stabilization (Fig. 1k,m) as density is clearly present also in the one-headed states (Fig. 1n,l). Whilst for ADP-Kif5b^MoNeIAK^ only the IAK tripeptide (KPI in the opposite direction) could be modelled (Fig. 1n), for AMPPNP-Kif5b^MoNeIAK^ the ordered region is longer (AQIAKP or AKPIRP) (Fig. 1l & Supplementary Fig. 8a). Overall, these observations suggest a mechanism for the formation of the autoinhibited zipped dimer in which a ‘IAK half-site’ binds first to one motor head leading then to the recruitment of the other motor by avidity.

### Association with MTs removes the Mg^2+^ ion from ADP-bound kinesin-1, resulting in an apo-like intermediate state

Currently, there is no high-resolution structure of the MT-bound kinesin-1 motor in the ADP state. Thus, our ∼2.9 Å resolution ADP-Kif5b^MoNeXXX^ structure fills an important gap in knowledge. For this chimera, which lacks the endogenous IAK motif, only one-headed MT-bound motors were observed that are devoid, as designed, of the extra density for the tail at its binding location (Fig. 1j and Fig. 2a). Density for the neck linker is also absent (Fig. 2b), indicating that it is undocked and flexible, in keeping with low-resolution reconstructions^20,46^. Noticeably, whilst ADP is present in its binding pocket, density for Mg^2+^ is absent (Fig. 2c), despite its clear presence at the N-site of α-tubulin (Supplementary Fig. 9a). Compared to the crystallographic ADP-bound Kif5b motor structure solved in the absence of MTs (PDB code 1bg2)^47^ (Fig. 2d), here, loop 9 that harbors the ‘switch I’ motif, unfurls from a short helical segment becoming partially disordered, loop 11 that contains the ‘switch II’ motif, adopts an ordered helical turn (Fig. 2e) whilst the MT-interacting helix α4 is extended (Fig. 2f). Overall, these observations indicate that, bar the presence of the bound nucleotide, the MT-bound structure of ADP-Kif5b^MoNeXXX^ resembles more the nucleotide-free conformation of Kif5b in complex with a tubulin dimer and a DARPin (PDB code 4lnu)^29^ than the MT-free ADP-bound structure of Kif5b (Supplementary Fig. 9b).

**Figure 2.**
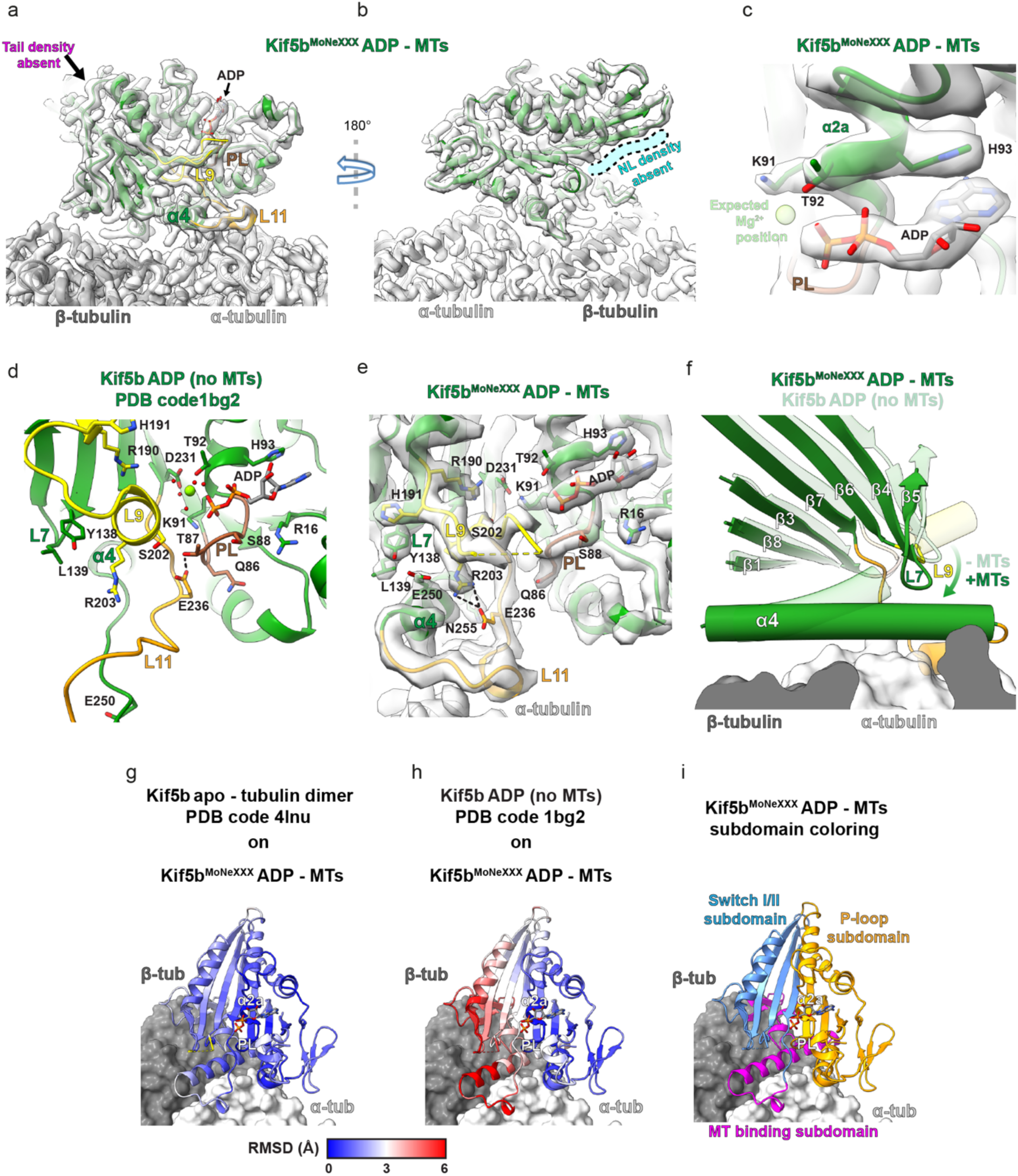
Kif5b motor domain binding to MTs causes dissociation of the Mg^2+^ ion from ADP and adoption of an apo-like conformation. **a**,**b.** Two views of the MT-bound Kif5b^MoNeXXX^ ADP- state reconstruction with the model fitted into density (grey transparent). In (**b**) the position where density for a docked neck linker (NL) could be expected (absent here) is indicated in semi-transparent cyan colour. **c.** View of the nucleotide pocket of MT-associated Kif5b^MoNeXXX^-ADP, highlighting the lack of cryo-EM density at the Mg^2+^ position (semi-transparent lime). **d,e.f.** Overviews of the nucleotide pocket and switch-motif containing loops 9 (L9) and 11 (L11), with side chains for key mechanistic residues shown for (**d**) the crystal structure of Kif5b-ADP motor domain without MTs (PDB code 1bg2) and (**e**) our Kif5b^MoNeXXX^-ADP structure in the presence of MTs (cryo-EM density in transparent grey). In (**f**), a superimposition highlighting helix α4’s extension, a twist of the overlying core β-sheet and concomitant movement of overlying loop 7 (L7) and loop 9 (L9) upon transition from the Kif5b-ADP state without MTs (PDB code 1bg2, semi-transparent model) to our MT-bound Kif5b^MoNeXXX^ ADP-state (opaque model). Helices are shown as tubes and αβ-tubulin is shown as grey surface representation. **g**,**h.** (**c**) The crystal structures of the Kif5b motor domain without nucleotide (apo) bound to αβ tubulin dimer (PDB code 4lnu) or (**d**) of Kif5b-ADP without MTs (PDB code 1bg2) are superimposed on our Kif5b^MoNeXXX^ ADP-state and calculated RMSDs are shown. The superimpositions in panels f-h use the nucleotide-holding P-loop and helix α2a elements for alignment. **i.** Kif5b^MoNeXXX^ ADP-state colored according to the kinesin motor domain subdomain scheme^26,29^.

The nucleotide-free (apo) conformation of kinesin-1 represents a ‘strong’ binding state^48^, with enhanced MT contacts formed by the ordering of the α4 helix extension and loop 11 region^29^. Structural superposition upon the P-loop/α2a helix that holds ADP reveals that our structure and the tubulin dimer-bound motor in the apo state exhibit low RMSD values globally (Fig. 2g), whilst MT-free Mg^2+^-ADP-bound Kif5b is structurally divergent on one side (Fig. 2h). If analyzed in terms of the kinesin subdomain scheme used in previous studies^26,29^ (Fig. 2i), the whole P-loop subdomain stays relatively unchanged while the switch I/II and MT-binding subdomains undergo significant conformational rearrangements. This results in a novel network of interactions between conserved residues of the extended helix α4/loop 11 region and loop 7/loop 9 of the switch I/II subdomain. One such new interaction is between R203 of switch I (in loop 9) and E236 of switch II (in loop 11) (Fig. 2e). In the MT-free Mg^2+^-ADP structure, the latter residue is bonded to T87 of the P-loop, thus locking the P-loop and loop 11 together in a manner crucial to stabilizing the Mg^2+^-ADP state^49^ (Fig. 2d). In our structure, loop 7/loop 9 and helix α4-proximal regions of the core β-sheet in the switch I/II subdomain (β strands 4-7) are drawn towards helix α4 and the MT-binding subdomain, while regions of the core β-sheet in the P-loop subdomain (β-strands 1, 3 and 8) remain relatively static (Fig. 2f). This causes ‘twisting’ of the core β-sheet that has been suggested to happen upon ADP release^29^. However, we show here that this occurs upon MT-binding and Mg^2+^ release prior to ADP exit.

### Structural basis for a two-step MT-stimulated ADP release mechanism

MT-stimulated ADP release has been proposed to be biphasic, with the first phase being inhibited by Mg^2+18,30,50^. This is consistent with an ADP-bound transition state lacking Mg^2+^ and representing the weak-to-strong MT-binding state. This has been suggested on the basis of FRET measurements to anticipate and accelerate ADP release^25^. Our cryo-EM sample preparation conditions have a >1000- fold molar excesses of Mg^2+^ and ADP compared to the motor. Yet, while Mg^2+^ density is absent, ADP is clearly present indicating that MT binding drastically reduces motor affinity for the ion, but has, together with Mg^2+^ release, a comparatively less pronounced effect on ADP. Retention of the latter, owing to its very high concentration, allowed us to analyze the conformational changes that impact on ADP affinity following MT binding and Mg^2+^ release.

The MT-induced structural changes discussed in the previous section induce significant displacements of key Mg^2+^-coordinating side chains of loop 9 and β7 relative to ADP, the ADP- binding P-loop and the helix α2a region in the P-loop subdomain that are not compatible with Mg^2+^ retention. In particular, prior to Kif5b interaction with the MT, conserved D231 on β7 indirectly coordinates Mg^2+^ via a water molecule and the hydroxyl group on helix α2a’s T92 (Fig. 3a). Upon MT binding, however, the Cα atoms of D231 are displaced by ∼3 Å relative to the P-loop and helix α2a, such that these interactions are broken with D231 forming instead a new hydrogen bonding network with conserved R190 on loop 9 (also enabled by its Cα atom moving ∼6.5 Å relative to the P-loop) and conserved K91 on α2a (Fig. 3a). Moreover, while interactions between ADP and the backbone of the P-loop and helix 2a as well as with the side chains of R14, R16, P17 and H93 are maintained, thus conferring residual stability to the nucleotide in its binding pocket, the β-phosphate-coordinating K91 side chain adopts a new rotamer that is H-bonded to D231 (Fig. 3b). The reorientation of K91, alongside the loss of β-phosphate-coordination by Mg^2+^ and its associated water network is therefore expected to lead to a significant loss of ADP affinity driving its release under physiological conditions.

**Figure. 3.**
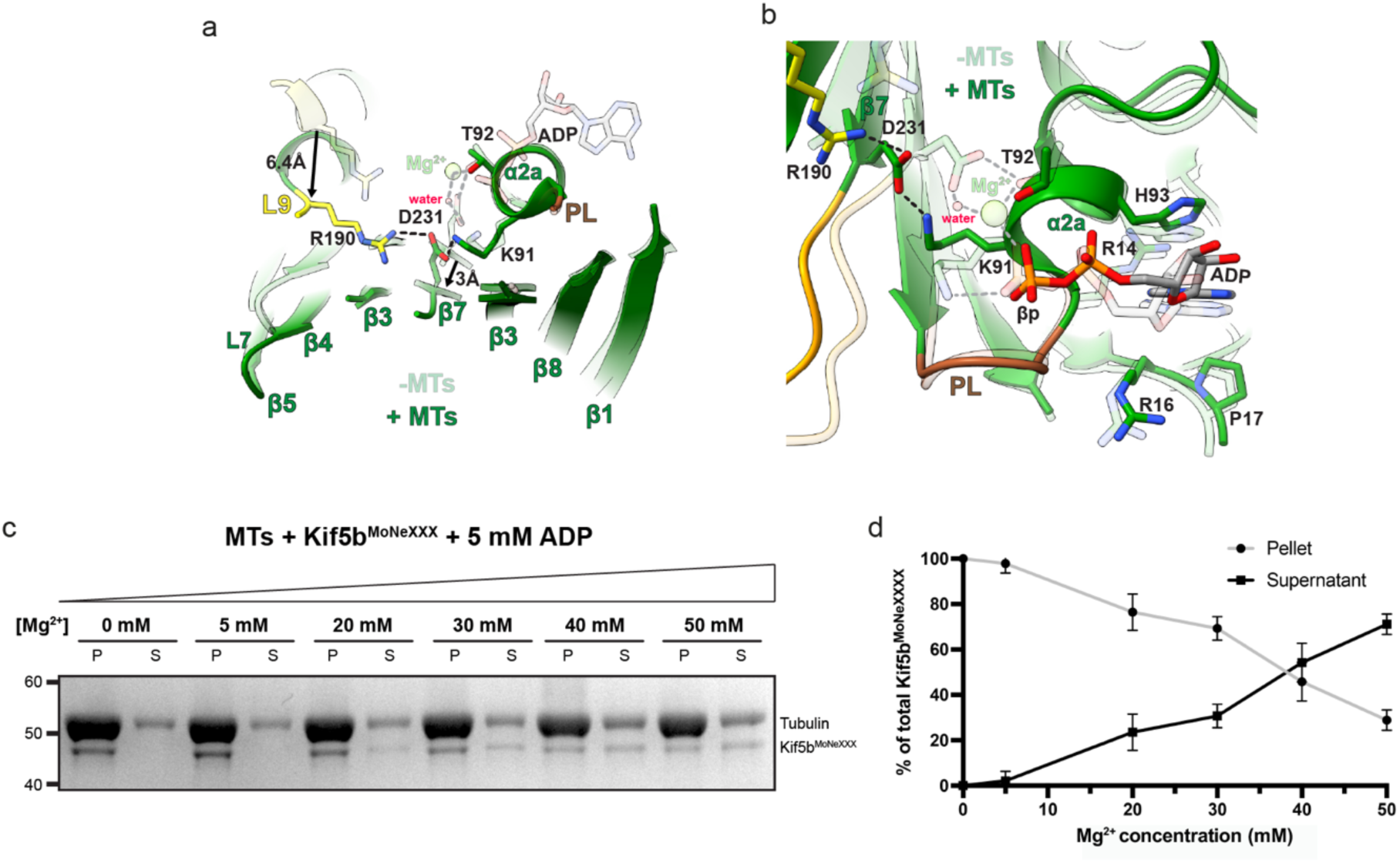
A two-step structural mechanism for MT-stimulated ADP release in Kif5b. **a**,**b**. Superimpositions of Kif5b-ADP without MTs (PDB code 1bg2, semi-transparent model) with our Kif5b^MoNeXXX^ ADP-state (opaque model) using the nucleotide-holding P-loop and helix α2a elements for structural alignment, highlighting the proposed (**a**) Mg^2+^-release and (**b**) ADP destabilization mechanisms. Hydrogen bonds in the Kif5b-ADP state without and with MTs are shown as grey and black dashed lines respectively. In (**a**), the movement of R190 and D231 α-carbon atoms are illustrated with black arrows. **c**. Representative co-sedimentation assay of MT and Kif5b^MoNeXXX^ co-incubated with increasing amounts of MgCl_2_ as indicated. Co-sedimentation was performed in BRB80 buffer with 5 mM ADP and the indicated concentrations of MgCl_2_, with taxol-stabilised MTs at 2 μM (tubulin dimer) and 0.5 μM Kif5b^MoNeXXX^. For conditions without Mg^2+^ we added 10 mM EDTA to remove any possible free Mg^2+^ ions. P = Pellet, S = Supernatant. Molecular weights are indicated to the left and bands are labelled to the right of the Coomassie-stained SDS-PAGE gels. **d.** Densitometric quantification of Coomassie-stained bands in the SDS-PAGE analysis of co-sedimentation assays (as in **c**), showing the percentage of total Kif5b^MoNeXXX^ in pellet vs. supernatant fractions. Each condition was repeated four times (standard error bars shown).

Given that the conformational changes driven by MT binding leading to a ‘strong’ MT-bound state are incompatible with Mg^2+^ remaining associated with ADP in the nucleotide pocket, we therefore predicted that in the presence of ADP, increasing Mg^2+^ concentrations would reduce MT affinity by preventing the conformational changes that lead to the ‘strong’ MT-bound state. To assess this, we performed MT co-sedimentation assays with Kif5b^MoNeXXX^ in the presence of ADP and increasing concentrations of Mg^2+^. As expected, MTs pellet at high centrifugal forces, while in the absence of MTs Kif5b^MoNeXXX^ remains chiefly in the supernatant (Supplementary Fig. 10). In the presence of a molar excess of MTs, we find that at [Mg^2+^] ≲ 10 mM all detectable Kif5b^MoNeXXX^ binds and co-pellets with the filaments (Fig. 3c). However, as [Mg^2+^] rise above ∼10 mM, Kif5b^MoNeXXX^ progressively moves from the pellet to the supernatant, indicating that MT binding is reduced in a [Mg^2+^]-dependent manner (Fig. 3c,d). We could not test Mg^2+^ concentrations above 50 mM due to significant MT destabilization, however, a large a molar excess is clearly needed to decrease Kif5b^MoNeXXX^ MT-association. This is consistent with the notion that MTs increase kinesin-1 ATPase rates by ∼1000 fold, by stimulating the rate-limiting Mg^2+^ and subsequent ADP release steps^16,50^. Overall, considering the apo-like conformation of our MT-bound and ADP-bound Kif5b^MoNeXXX^ structure, we believe to have captured kinesin-1 in the pre-ADP release ‘strong’ MT-binding state, induced by Mg^2+^ dissociation.

To our knowledge, an Mg^2+^-free ADP-bound apo-like conformation of Kif5b on MTs as described here has only been observed previously in the atypical kinesin Kif14^51^. When the motor domain of the latter (PDB code 6wwm) was superimposed onto the motor domain in our ADP-bound Kif5b^MoNeXXX^ model, the structures showed high similarity apart from the divergent loops 2, 3 and 8 and 9, which was similar at its base but more ordered at its apex in Kif14 (Supplementary Fig. 11a). Furthermore, a similar ‘apo-like’ network of key conserved residues was seen (Supplementary Fig. 11b), including the interactions between R591, D638 and K488 (equivalent to Kif5b R190, D231 and K91 respectively) which we regard as critical for the Mg^2+^ and ADP release mechanisms (Fig. 3a,b). Kif14 also exhibits an apo-like ADP-bound conformation lacking Mg^2+^ in the absence of MTs^52^, except for loop 11 being more disordered and loop 9 being more disordered at its apex (Supplementary Fig. 11c). Compared to other kinesins, Kif14 has an uncommonly high affinity for MTs in the presence of ADP^52^, which can be explained by the lack of Mg^2+^ coordination and the adoption of this apo-like conformation including an extended/stabilized helix α4 MT-interacting element, even in the absence of MTs. The extension and orientation of helix α4 in Kif14 in the absence of MTs coincides with a core-β twist and movement of the overlying loop 9, leading to the R591 and D638 interaction (R190 and D231 in Kif5b) precluding Mg^2+^ binding (Supplementary Fig. 11c). An apo-like conformation in solution explains the unusually high basal ATPase rate of Kif14^52^, as ADP-release is accelerated by the lack of Mg^2+^ coordination. Interestingly however, MTs still stimulate ATPase in this motor, but only by a further 3-fold as opposed to more than ∼104 fold in kinesin-1^16,52^. When comparing MT-bound^51^ and MT-free^52^ ADP-Kif14 structures, this can likely be explained by the reorientation of K488 side chain upon MT interaction to interact with D638 instead of ADP’s β-phosphate, further reducing ADP affinity and accelerating its release (Supplementary Fig. 11b,c).

### Tail association, with or without zippering of the distal head, does not affect the ADP-bound conformation of the MT-bound head

To assess the structural impact of the tail on the ADP-bound conformation, we superimposed the one-headed ADP-bound Kif5b^MoNeIAK^ model onto ADP-bound Kif5b^MoNeXXX^. The models are almost identical in all regions (RMSD=0.3 Å), demonstrating that tail association has little effect on the ADP-bound structure of the MT-bound head (Fig. 4a). When superimposing the one-headed and two-headed ADP-bound Kif5b^MoNeIAK^ models on the MT-associated head (Fig. 4b), again negligible structural differences were observed (RMSD=0.2 Å). Despite the presence of the tail, the MT-bound head of Kif5b^MoNeIAK^ adopts the same apo-like ADP-bound and Mg^2+^-less conformation seen in Kif5b^MoNeXXX^, with an extended helix α4 and ordered loop 11 in the MT-binding subdomain and an unfurled partially disordered loop 9 (Fig. 4c). Furthermore, the bonding network between loop 9’s R190, β7’s D231 and helix α2a’s K91 described for Kif5b^MoNeXXX^ (Fig. 3b) is also present in the MT-associated head of one- (Fig. 4d) and two-headed states of Kif5b^MoNeIAK^ (Fig. 4e). This strongly suggests that the IAK-motif tail, irrespective of the presence of the zipped distal head, has little effect on the MT-stimulated Mg^2+^ or ADP-release mechanisms of Kif5b.

**Figure 4.**
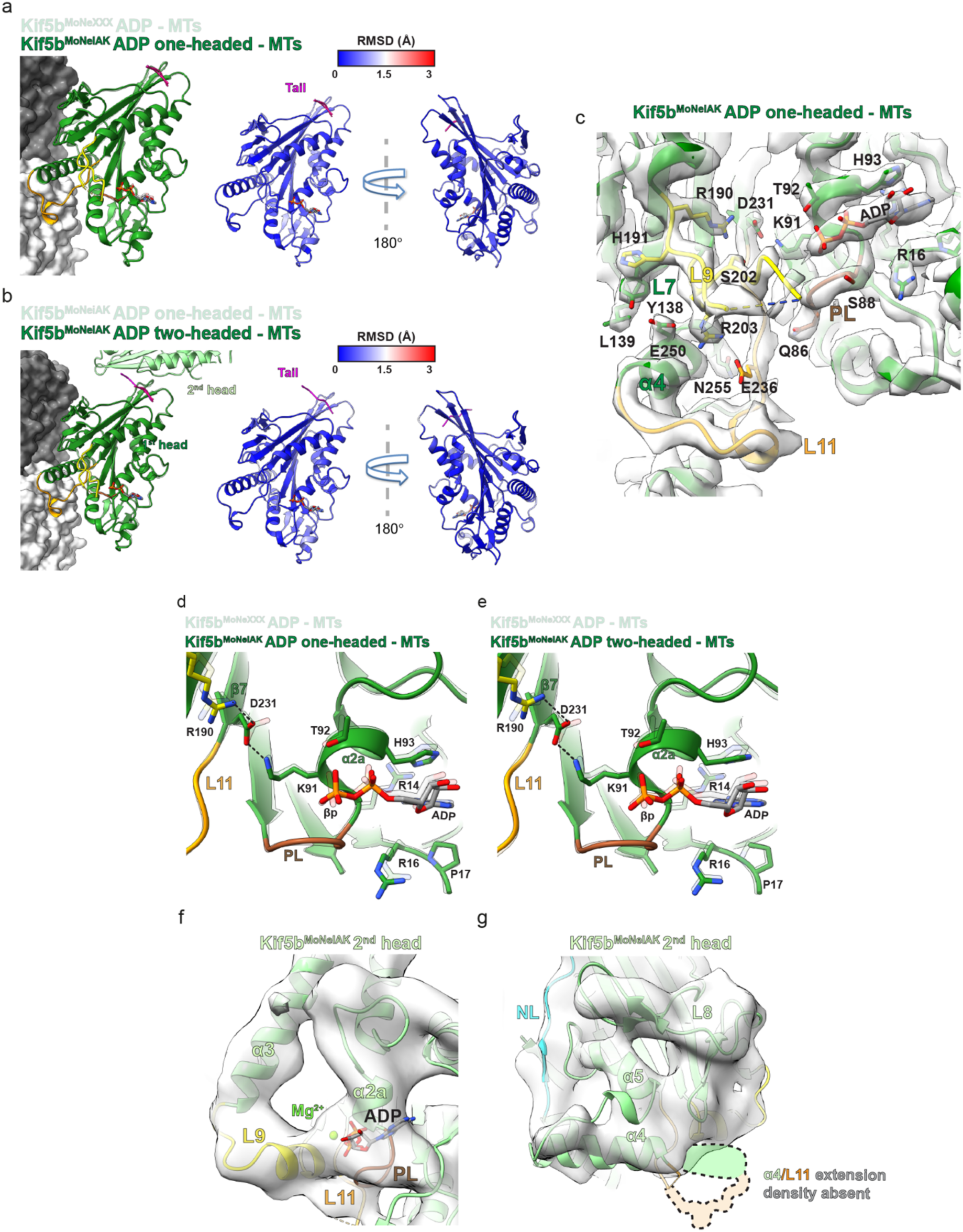
Tail or 2^nd^ head association has no effect on the ADP-bound conformation of the MT- associated Kif5b motor domain. **a.** ADP-bound motor domains of the Kif5b^MoNeIAK^ and Kif5b^MoNeXXX^ (semi-transparent) MT-associated models were superimposed (left). RMSDs were calculated and displayed on the Kif5b^MoNeIAK^ model (right), with the Kif5b^MoNeIAK^ tail shown in magenta. **b.** MT-associated ADP-bound motor domains of the Kif5b^MoNeIAK^ one (semi-transparent) and two-headed models were superimposed (left). RMSDs were calculated and displayed on the Kif5b^MoNeIAK^ two-headed model (right) with the Kif5b^MoNeIAK^ tail shown in magenta. **c.** Overview of the nucleotide-pocket and switch-motif containing loops 9 and 11 (L9 and L11) in the ADP-bound Kif5b^MoNeIAK^ one-headed model, with side chains for key mechanistic residues shown. Cryo-EM density is shown as semi-transparent grey. **d,e.** Superimposition of the ADP-bound MT-associated motor-domains of Kif5b^MoNeXXX^ (semi-transparent) and; **(d)** the Kif5b^MoNeIAK^ one-headed model, **(e)**, the Kif5b^MoNeIAK^ two-headed model. A view of the nucleotide binding site is shown, alongside key side chains involved in Mg^2+^ and ADP-release. **f,g.** Views of the 2^nd^ (not MT-associated) head in the Kif5b^MoNeIAK^ two-headed model in the presence of ADP. Cryo-EM density (semi-transparent) was low-pass filtered to 6 Å resolution for resolution-appropriate visualization. The position where density for an extended helix α4 and ordered L11 could be expected (but here is absent) is indicated in semi-transparent colors with dashed black outlines.

Supporting the role of MT binding in the conformational change observed in the MT-associated head leading to Mg^2+^ release, density (although at lower resolution and low-pass filtered to 6 Å for reliable interpretation) in the distal head of Kif5b^MoNeIAK^ is most consistent with loop 9 transitioning to a short helical segment, helix α4 being short and partly disordered, loop 11 being mainly disordered and Mg^2+^-ADP remaining associated (Fig. 4f,g and Supplementary Fig. 12). This strongly resembles the Mg^2+^-ADP-bound conformation observed previously for tail-less Kif5b motor domain without MTs^47^.

### Tail association, with or without zippering of the distal head, does not affect the ATP-hydrolysis competent conformation of the MT-bound head

To assess the structural impact of the tail on the ‘ATP-like’ conformation, we superimposed the one-headed AMPPNP-bound Kif5b^MoNeIAK^ model onto the X-ray crystallographic model of ADP-AlF_4_- bound Kif5b in complex with a tubulin dimer and a DARPin (PDB code 4hna)^15^ (Fig. 5a). The models superimposed extremely well in all regions of the motor domain (RMSD=0.5 Å), demonstrating tail-association has little effect on the ATP-like state of the MT-associated head (Fig. 5a). Upon superimposing the one-headed and two-headed AMPPNP-bound Kif5b^MoNeIAK^ models on the MT-associated head (Fig. 5b), again a high level of structural similarity was observed (RMSD=0.2 Å). This indicates that even the additional presence of the 2^nd^-head zipped via the IAK motif has negligible or no structural effect on the MT-bound head. Despite the presence of the tail or the zipped 2^nd^ head therefore, the MT-bound head has an extended helix α4 and ordered loop 11 in the MT-binding subdomain, associated with a ‘strong’ MT-bound state (Fig. 5a,b,c). The helical turn of switch-II containing loop 11 contacts a fully ordered switch-I-containing Loop 9, closing AMPPNP and coordinated Mg^2+^ between these elements and the P-loop (Fig. 5c). The high-resolution structure of a kinesin-5 (Eg5)^53^ demonstrated that this ‘closed’ conformation allows ATP’s γ-phosphate to be stabilized by Mg^2+^ and residues equivalent in Kif5b to S201 and S202 of switch I in loop 9, whilst preparing it for nucleophilic attack via waters arranged by S202 and R203 of switch I (in loop 9) and G234 and E236 of switch II (in loop 11) (Fig. 5d,e & Supplementary Fig. 13).

**Figure 5.**
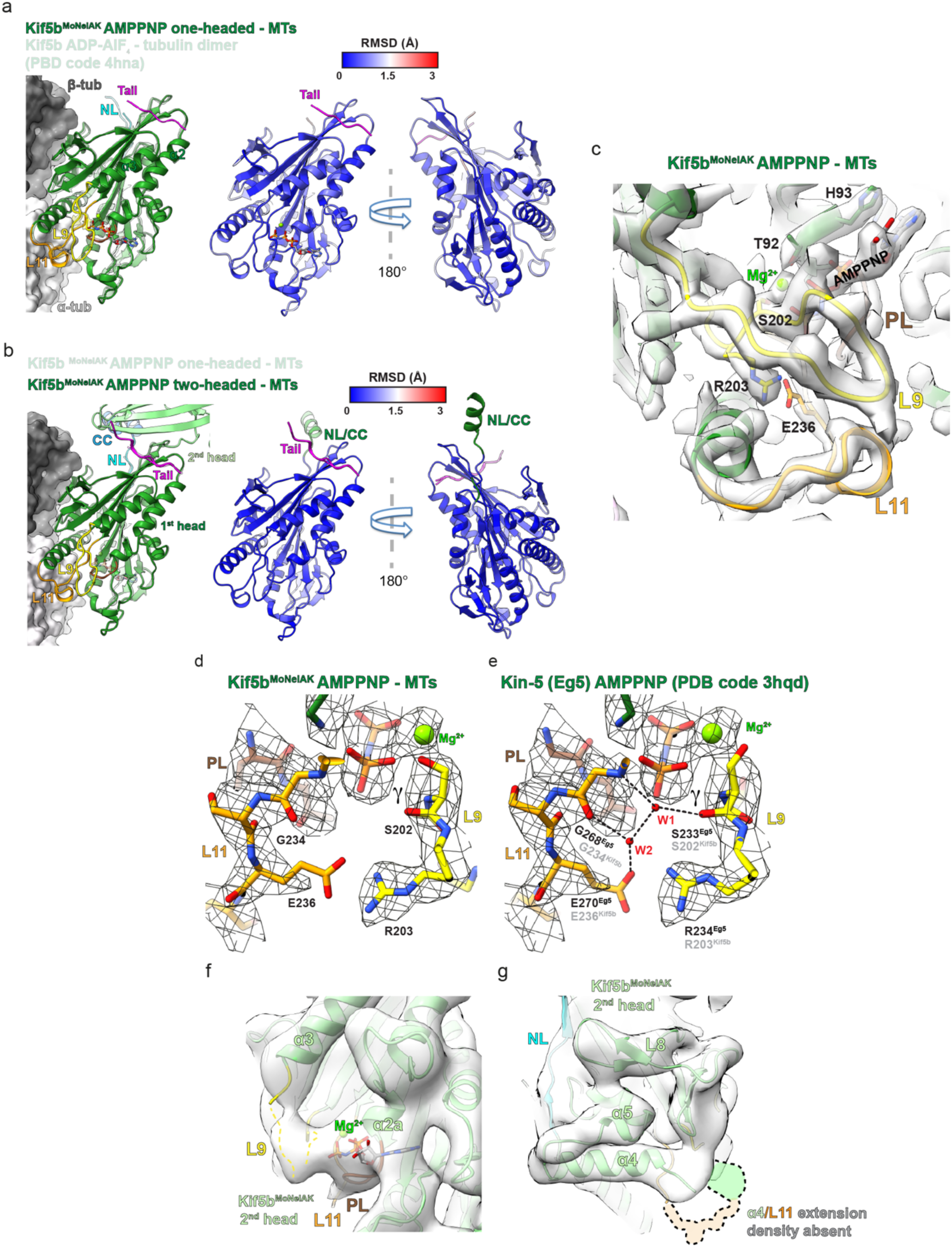
Tail or 2^nd^ head association has no effect on the ATP-like conformation of the MT- associated Kif5b motor domain. **a.** Motor-domains of the Kif5b^MoNeIAK^ AMPPNP and MT-bound and ADP-AlF_4_ and tubulin-bound (semi-transparent, PDB code 4hna) models were superimposed (left). RMSDs were calculated and displayed on the Kif5b^MoNeIAK^ model (right), with the Kif5b^MoNeIAK^ tail shown in magenta. **b.** MT-associated AMPPNP-bound motor domains of the Kif5b^MoNeIAK^ one (semi-transparent) and two-headed models were superimposed (left). RMSDs were calculated and displayed on the Kif5b^MoNeIAK^ two-headed model (right) with the Kif5b^MoNeIAK^ tail shown in magenta and the NL-CC region shown in green. **c.** Overview of the nucleotide-pocket and switch-motif containing loops 9 and 11 (L9 and L11) in the AMPPNP-bound Kif5b^MoNeIAK^ one-headed model, with side chains for key conserved residues shown. Cryo-EM density is shown as semi-transparent grey. **d,e.** Close up view of residues involved in ATP hydrolysis, comparing **(d)** our MT-associated AMPPNP-bound Kif5b^MoNeIAK^ motor domain (left) and **(e)** Eg5 motor domain (PDB code 3hqd). Both models are fitted into density for the MT-associated motor domain of AMPPNP-bound Kif5b^MoNeIAK^ (mesh). The high resolution in the Eg5 motor domain crystal structure allows display of the two water molecules and associated bonding network (black dashed lines) that are involved in nucleophilic attack on the γ-phosphate^53^. In the right-hand panel, Eg5 residue numbering is shown in black and Kif5b residue numbering in grey. **f,g.** Views of the 2^nd^ (not MT-associated) head in the Kif5b^MoNeIAK^ two-headed model in the presence of AMPPNP. Cryo-EM density (semi-transparent) was low-pass filtered to 6 Å resolution for resolution-appropriate visualization. The position where density for an extended helix α4 and ordered L11 could be expected (but here is absent) is indicated in semi-transparent colors with dashed black outlines.

In the two-headed AMPPNP-Kif5b^MoNeIAK^ state, the 2^nd^ head was resolved at lower resolution (∼4-6 Å) particularly at regions distal to the tail and tip of the motor domain (Supplementary Fig. 4c), such that low-pass filtering its density to 6 Å was appropriate to examine its conformation. Nonetheless, this revealed that while Mg^2+^-AMPPNP occupied the nucleotide pocket, helix α4 was not extended, and loops 9 and 11 were not well ordered, such that the 2^nd^ head was not in a nucleotide-hydrolysis competent conformation (Fig. 5f,g). This finding is consistent with the requirement of MT binding for ordering of these secondary structure elements and the adoption of a fully hydrolysis-competent conformation^15,17,18,20^.

### MT-bound Kif5b^MoNeIAK^ undocks its neck-linker in the presence of ADP regardless of tail binding

In the presence of MTs, closing of the nucleotide pocket in response to ATP analogues induces conformational changes in kinesin motor domains leading to extension of helix α6 and docking of the preceding neck-linker^20,24,54^. After Pi release following hydrolysis, the neck linker then undocks and becomes disordered, along with the C-terminus of helix α6, through both Mg^2+^-ADP-bound and apo stages of the ATPase cycle^19,20^. In contrast, in the absence of MTs neck linker docking is not coupled with the nucleotide state^55^. In the crystallographic structure of the ADP-bound *Drosophila* kinesin-1 motor dimer in complex with an IAK motif peptide, the neck linkers in both heads are docked, leading into the coiled-coil^35^. From this structure, the tail was suggested to prevent neck linker undocking and that this mechanism inhibits MT-stimulated ADP release (double-lockdown mechanism).

In the presence of AMPPNP, the MT-associated head of both single and two-headed Kif5b^MoNeIAK^ states exhibited an extended helix α6 and docked neck linker (Fig. 6a,b) coincident with closing of the nucleotide pocket around the ATP analogue (Fig. 5c). In the two-headed state, helix α6 was extended, and the neck-linker was docked in the 2^nd^ head as well, such that a portion of the dimeric coiled-coil could be resolved (Fig. 6b and Supplementary Fig. 14a). In contrast to a processive dimer however, the 2^nd^ head is not thrust forward onto the next αβ-tubulin binding site towards the MT plus end but is instead locked in a restricted orientation against the 1^st^ head by the tail (Fig. 6b & Supplementary Fig. 7a). In contrast, in the presence of ADP, the MT-associated head of Kif5b^MoNeIAK^ clearly exhibited a disordered (and undocked) neck linker and shortened helix α6 in both single- and two-headed states (Fig. 6c,d). In contrast to the MT-associated head, the 2^nd^ head in the two-headed state has an extended helix α6 and docked neck-linker (Supplementary Fig. 14b,c). Consistent with the lack of neck linker docking in the MT associated head, we did not observe ordering of CC1 (Fig. 6d and Supplementary Fig. 14b). As the two-headed class had lower occupancy in the presence of ADP compared to AMPPNP (Fig. 1i), it is likely that the stabilization of a docked neck linker and CC1 (Fig. 6b) supports a two-headed complex including a more ordered tail (Fig. 1k,m). Regardless, these reconstructions therefore demonstrate that nucleotide-dependent neck linker undocking still occurs in the MT-associated head, regardless of the presence of the autoinhibitory tail with or without a zippered 2^nd^ head.

**Figure 6.**
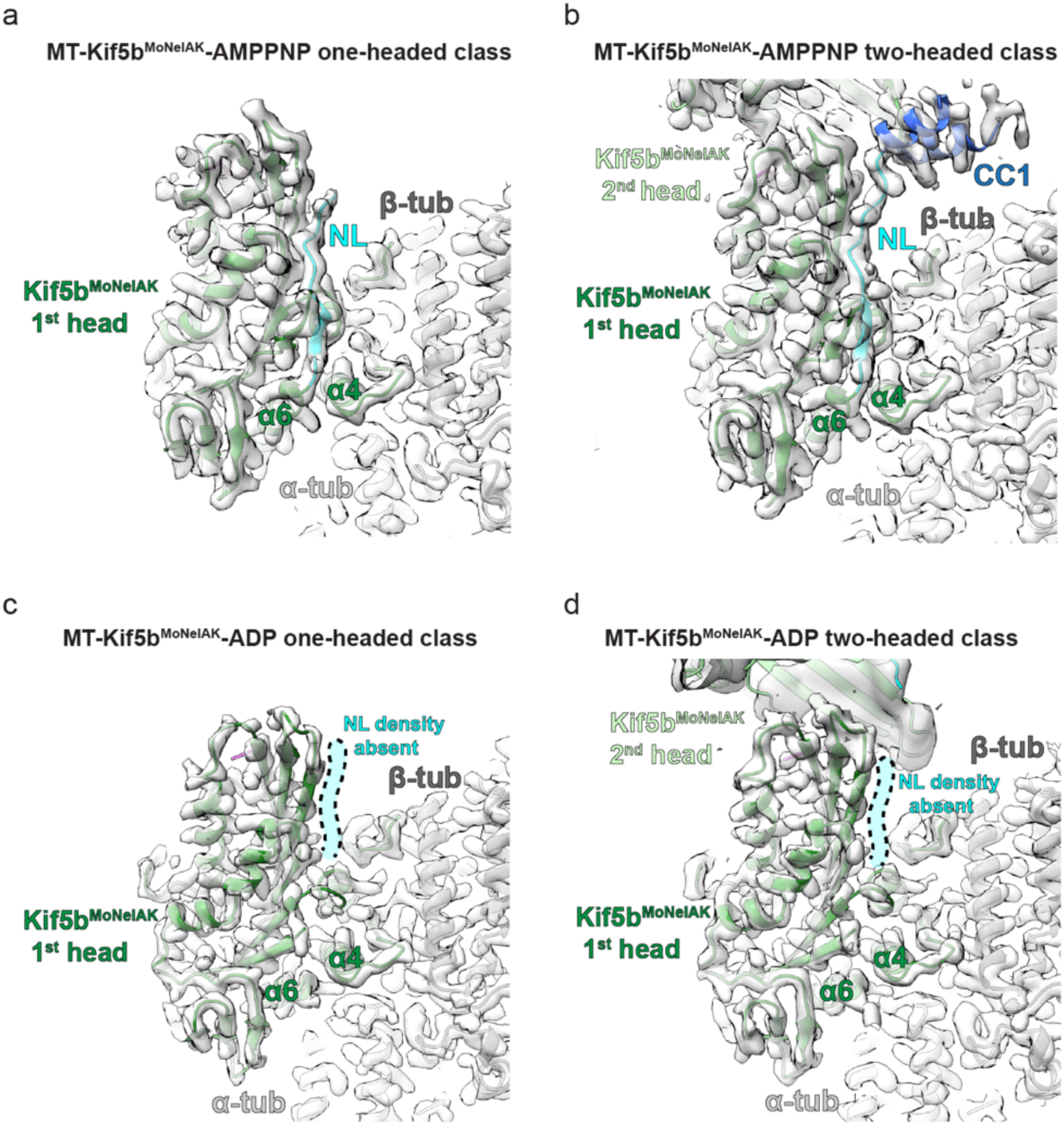
MT-associated Kif5b^MoNeIAK^ undocks its neck-linker in the presence of ADP. **a,b,c,d.** Views of the neck-linker and coiled-coil of the MT-associated head of Kif5b^MoNeIAK^ bound to **(a,b)** AMPPNP or **(c,d)** ADP in **(a,c)** one- or **(b,d)** two-headed states, with corresponding density in semi-transparent grey. In panels **(c)** and (**d)** the position where density for a docked neck-linker could be expected (but here is absent) is indicated in semi-transparent cyan.

## Discussion

MT-induced stimulation of ATP hydrolysis and cargo-dependent relief of autoinhibition are key aspects of kinesin-1’s mechanochemistry as they contribute to preventing futile mechanochemical cycles in the absence of productive motor runs. The rate-limiting step in the kinesin-1 ATPase cycle is ADP release^16^ and various studies have shown that this follows a two-step sequential mechanism whereby Mg^2+^ release precedes and accelerates that of ADP, with the weak-to-strong MT-binding state transition anticipating the ADP release step^18,25,30,50^.

Our high-resolution structure of ADP-loaded Kif5b bound to its MT track captured the ‘strong’ MT-bound state of the motor after Mg^2+^ dissociation and before ADP release, allowing us to describe a structural model for the two-step mechanism of MT-stimulated ADP release (Fig. 7). Transition to the ‘strong’ binding state coincides with the ordering of the MT-interacting helix α4 and loop 11 elements prior to ADP release. This produces a new set of interactions with the core β-sheet and associated loop 7 and loop 9, causing a ‘twist’ of the β-sheet and displacement of these loops relative to the nucleotide-holding P-loop/helix α2a. As a result, interactions between the conserved D231 of the core β-sheet (a switch-II motif residue), T92 (helix α2a) and the water ‘cap’ holding the Mg^2+^ ion are broken, leading to its release (step 1). Concomitantly, D231 forms a new hydrogen bonding network with R190 on the displaced loop 9 and with K91 on helix α2a, such that neither Mg^2+^ or K91 remain bound to the β-phosphate of ADP, suggesting a mechanism for MT-dependent and Mg^2+^-loss-triggered stimulation of ADP release (step 2). A recent magic-angle-spinning NMR study of nucleotide-free Kif5b bound to MTs also shows an interaction between K91 and D231, as in our ADP-bound structure^56^. The importance of conserved D231 and R190 as part of the ‘Mg^2+^- stabilizer’ network has previously been noted, and replacement of these residues drastically reduced the ATPase rate whilst increasing ADP release rates in kinesins^29,30^. The key role of Mg^2+^ in ADP retention is also clear from the observation that mutation of the P-loop Mg^2+^-coordinating residue T92 results in a drastic loss of ADP affinity^49^. Our observation that K91 additionally no longer supports the ADP β-phosphate suggests this as an auxiliary factor in reducing ADP affinity in response to MT binding. K91 is a highly conserved residue in the P-loop Walker A motif (GxxxxG<u>K</u>(S/T)) that spans P-loop and helix α2a structural elements, and a substitution at the site was recently identified as causing kyphomelic dysplasia^57^.

**Figure 7.**
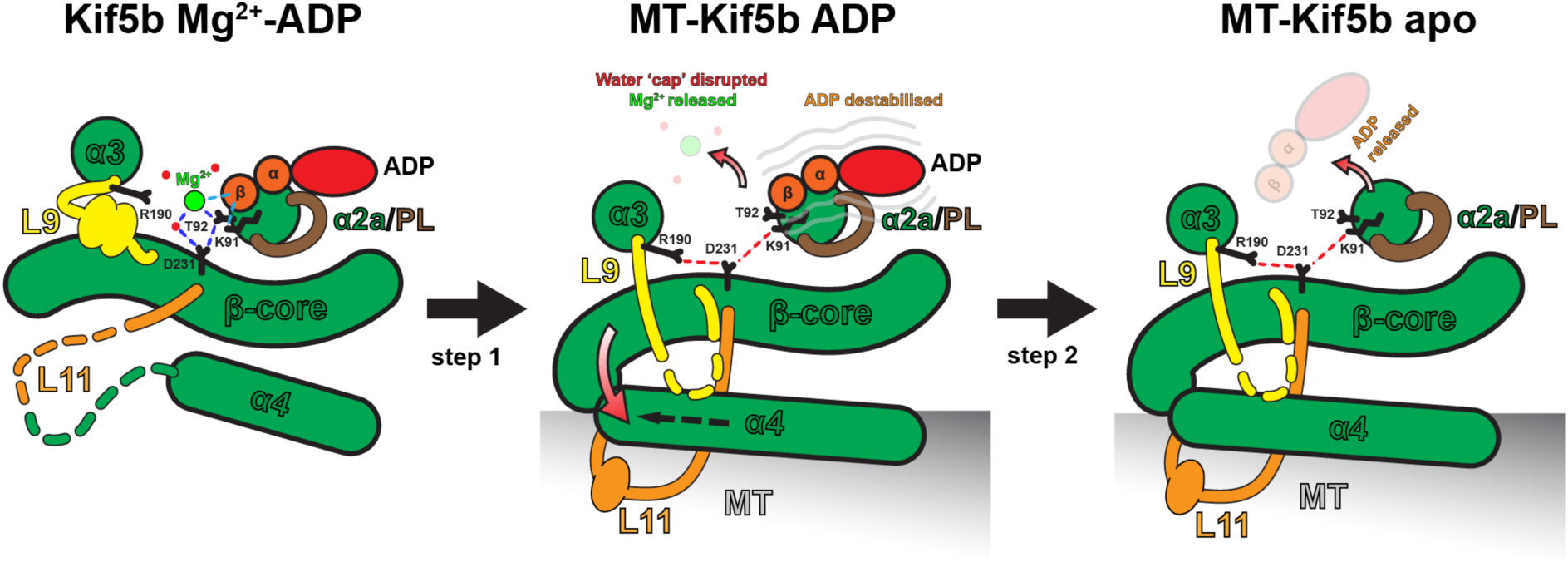
Model for a two-step structural mechanism of MT-stimulated ADP release. Schematic illustrating a proposed two-step structural mechanism of MT-stimulated ADP release, showing movements of key SSEs and side chains relative to the static helix α2a and P-loop. MT binding triggers transition to an apo-like state, including helix α4 extension and twisting of core β-sheet (β- core) along with movements of overlying helix α3 and loop 9 elements. These movements dismantle a hydrogen bonding network involving D231 and T92 that coordinates Mg^2+^ through a water ‘cap’ (blue dashed lines). This drives Mg^2+^ out of the complex, with D231 forming a new bonding network with R190 and K91 (red dashed lines). These changes disrupt key bonds stabilizing ADP’s β- phosphate (cyan dashed lines), leaving ADP only weakly coordinated by remaining contacts with helix α2a and the P-loop, promoting its subsequent release. α and β are α-phosphate and β-phosphate, respectively.

Another finding of the present study is that the IAK motif containing tail, even when zippering the second head, has no structural effects on the MT-bound motor in the presence of either ADP or AMPPNP. Therefore, any influence of the IAK tail on kinesin-1’s mechanochemistry is unlikely to be mediated by steric or allosteric effects on the nucleotide binding site affecting nucleotide binding, release or hydrolysis. This contrasts with a previous ∼8 Å cryo-EM study of a kinesin-1 motor domain without nucleotide which found tail-peptide density alongside the switch I-motif containing loop 9, which was suggested to inhibit ADP-release by preventing conformational changes around the MT-binding region and nucleotide pocket upon MT binding^34^. However, as this sample was chemically cross-linked, sample preparation artefacts cannot be ruled out. The location of the bound tail and the mode of zippering of the second head observed in our study is akin to that seen in the X-ray crystallographic structure of the MT-free ADP-bound autoinhibited *Drosophila* kinesin-1 dimer^35^. In the latter structure, CC1 was ordered with the neck linker docked in both heads prompting the authors to suggest a ‘double-lockdown autoinhibition mechanism’, whereby the tail locks the two heads in a restricted orientation (lock 1) that, in turn, prevents neck linker undocking (lock 2) such that ADP release is inhibited. However, we unambiguously show here that in the presence of ADP, the MT-associated motor domain undocks its neck linker regardless of the presence of the bound tail or of the distal motor head zippered to the first. This implies that the ‘double-lockdown’ mechanism of autoinhibition needs to be refined.

What role does tail binding then play in kinesin-1 autoinhibition? Full-length kinesin-1 with or without KLCs is autoinhibited in a manner that strongly suppresses MT association^33,39,40^ and according to recent models based on low-resolution negative stain EM and cross-linking mass-spectrometry, autoinhibited full-length kinesin-1 folds such that regions of the stalk obscure a MT-binding surface of at least one of the motor domains^36–38^. In these models, one of the two IAK-motif containing tails in the dimer folds back to associate with and zip up the two motor domains (as in the current study), thanks to an ‘elbow’ that allows flexibility in the stalk. Although kinesin-1 appears to have a built-in tendency to fold back on itself^37^, the presence of the IAK motif-containing tail is required for full inhibition of ATPase activity, ADP release and motility^31,33,38,40^. Taken together, in full-length kinesin-1 a key function of the IAK motif-containing tail appears that of acting as a ‘safety belt’ ensuring a stable compacted fold, thereby indirectly preventing MT landings due to regions of the stalk sterically blocking the MT-binding interface. However, although MT landing events of cargo-free full-length kinesin-1 are minimal, a basal level remains and when MT-associated these motors display reduced motility with frequent pausing and longer association/run durations^37,40^. Furthermore, truncated dimeric kinesin-1 constructs lacking the tail, and in some cases most of the coiled-coil, associate strongly with MTs but are stalled and display reduced MT-stimulated ATPase and ADP release rates in the presence of exogenous tail peptide^41,42,44^. Therefore, this represents an additional autoinhibitory mechanism mediated by the IAK-containing tail independent of simple reduction of MT-binding via steric inhibition in the folded conformation. The structures presented here demonstrate that this does not arise from any direct influence of the tail on the conformational transitions of the MT-associated head and suggest that initial MT-stimulated ADP-release, ATP-binding or hydrolysis steps are unaffected. However, our structures would indicate that upon ATP binding to the MT-associated head, the distal one is not thrust forward by neck linker docking but is instead restricted by the tail in a new orientation dictated by the two-headed complex. Consequently, the second head is not presented to the next binding site on the MT track thus preventing MT-stimulated ADP release and ultimately processive motility. In support of this hypothesis, an initial transient of MT-stimulated ADP release occurs in the presence of the tail before further ADP release is prevented, with a similar effect found when chemically crosslinking the heads together in a tailless construct^35^. This mechanism can also account for tail-dependent stalling of kinesin-1 on MTs in a strongly associated state^37,39–42,44^. As only one of the two tails of kinesin-1 associates with the heads^35^ and promotes autoinhibition^32^, it is possible that intermolecular interactions with other kinesin-1 molecules (Supplementary Fig. 15) results in zippering leading to cross-motor inhibition resulting in the characteristic tail-dependent pausing events. Autoinhibition in kinesin-1 therefore appears to be composed of multiple mechanistic layers, including those dependent on steric hinderance of MT association by regions of the stalk and those dependent on the reduction of processive stepping behavior.

## Materials and Methods

### Protein expression and purification

A codon-optimized DNA sequence of human Kif5b (Uniprot entry P33176) encoding the motor domain, neck linker and CC1 (residues 1-357) fused via a flexible (Thr-Gly-Ser)_9_ linker to the C-terminal tail region encompassing the autoinhibitory IAK motif (residues 912-935) was purchased from Genscript and subcloned between the NdeI/XhoI sites of a pET28 vector (Novagen). We refer to this chimeric protein as Kif5b^MoNeIAK^ for Kif5b <u>Mo</u>tor-<u>Ne</u>ck-<u>IAK</u>. A second construct was also synthesized that replaces the IAK motif region (919-Q<u>IAK</u>PIR-925) with the flexible palindromic sequence (TGSTSGT). We refer to this construct as Kif5b^MoNeXXX^.

Kif5b^MoNeIAK^ and Kif5b^MoNeXXX^ were expressed in the *E.coli* BL21(DE3) strain. Briefly, single colonies were picked and grown at 21°C overnight in Lysogeny Broth (LB) supplemented with kanamycin. Small-scaled overnight cultures were used to inoculate large-scale cultures (1:500 ratio) in Terrific Broth (TB) supplemented with 2mM MgCl_2_, 0.1% glucose and antibiotic. Cell cultures were incubated at 37 °C in a shaking rotator until OD600 reached 0.9-1.0. Protein expression was induced with 0.4mM isopropyl β-D-1-thiogalactopyranoside (IPTG) at 18 °C, for 16h. Cells were harvested by centrifugation (5,000g x 20 min, 4 °C) and the cell pellet resuspended in lysis buffer composed of 20 mM Tris pH 8.8, 200mM NaCl, 2mM MgCl_2_, 1mM EGTA, 1mM MgATP and 5mM β-mercaptoethanol (β-ME) supplemented with protease inhibitor cocktail (Merck) and benzonase endonuclease (10 U/ml) (Merck). After homogenization, cells were lysed by sonication on ice, followed by centrifugation (20,000g x 45min, 4°C). The soluble material was subsequently filtered using a 0.22 mm pore size filter and was loaded onto a 5 ml HisTrap HP column (GE Healthcare), which was pre-equilibrated with lysis buffer and immobilized metal affinity chromatography (IMAC) was performed at 4 °C. Target proteins eluted in an imidazole linear gradient, were buffer exchanged in lysis buffer, using dialysis membrane (with 3.5 MWCO) for 3h. Kif5b^MoNeIAK^ was further incubated overnight at RT with thrombin conjugated beads (Thrombin CleanCleave Kit, Sigma-Aldrich) for His-tag cleavage. The protein-bead mixture was then filtered using a gravitational column, and the untagged protein was collected by reverse IMAC on a HisTrap HP column (GE Healthcare) pre-equilibrated with lysis buffer. Untagged Kif5b^MoNeIAK^ was buffer exchanged using a PD-10 desalting column in 50 mM Hepes pH 7.5, 20mM NaCl, 2 mM MgCl_2_, 1mM EGTA, 1mM MgATP and 5mM β-ME. The protein was further purified by Ion Exchange Chromatography (IEX) using a RESOURCE S cation exchange column (Cytiva) and protein was eluted in a salt gradient (50 mM Hepes pH 7.5, 1M NaCl, 2 mM MgCl_2_, 1mM EGTA, 1mM MgATP, 5mM β-ME), followed by size exclusion chromatography (SEC) on a HiLoad 16/600 Superdex 200 column (GE Healthcare) pre-equilibrated with 25 mM HEPES pH 7.5, 150 mM NaCl, 2mM MgCl_2_, 1mM EGTA, 20μΜ MgATP and 5mM β-ME. Kif5b^MoNeXXX^ was purified by SEC immediately after the IMAC step without His_6_-tag cleavage using the same buffer conditions employed for Kif5b^MoNeIAK^.

### MT Preparation

Purified and lyophilized porcine brain tubulin (>99% pure, Cytoskeleton Inc.) was reconstituted on ice to 10mg/ml in BRB80 (80 mM PIPES, 2 mM MgCl_2_, 1 mM EGTA, 1 mM DTT, pH 6.8) then snap frozen in liquid nitrogen and stored at −80°C. Upon thawing, the tubulin was centrifuged at 13,200 RPM on a desktop centrifuge at 4°C to remove large aggregates, then the supernatant extracted for further use. To polymerize MTs, the supernatant was then diluted to 5 mg/ml tubulin in BRB80 with 1 mM GTP (Sigma) and incubated at 37°C for 45 minutes. A solution of 1 mM paclitaxel (Calbiochem) in DMSO was then added and MTs incubated at 37°C for another 45 minutes. Stabilized MTs were left at room temperature for at least 24 hours before use.

### MT co-sedimentation assays

In 7 x 20mm Thickwall Polycarbonate tubes (Beckman Coulter), paclitaxel-stabilized MTs were mixed at room temperature with Kif5b proteins in BRB80 plus indicated the nucleotides or metal salts. Controls with no MTs or no Kif5b were included. After incubating for 10 minutes, the samples were centrifuged at 85,000 RPM in an Airfuge® Ultracentrifuge (Beckman Coulter). The supernatants were extracted and mixed 1:1 with 2x SDS-PAGE sample buffer (ThermoFisher Scientific). The pellets were then resuspended with ice cold BRB80, then further mixed and resuspended with 2x SDS-PAGE sample buffer and extracted. Pellet and supernatant samples were analyzed by SDS-PAGE and stained with InstantBlue® Coomassie Protein Stain (Expedeon). For densitometric quantification, background-subtracted band intensities were measured from lanes in the Coomassie-stained SDS-PAGE gel with the ‘Gel Analyzer’ tool in FIJI software^58^.

### Sample preparation for cryo-EM

MTs were diluted in BRB80 at room temperature to 0.16 mg/ml before use. Kif5b was diluted in BRB80 to 1 mg/ml and preincubated with 5 mM AMPPNP or ADP for 20 minutes at 4°C before use. A volume of 3 μl of MTs were pre-incubated on glow-discharged C-flat^TM^ carbon-coated copper EM grids (2 μm holes with 2μm hole spacing, Protochips) at room temperature for 45 seconds, excess buffer manually blotted away, then 3μl of Kif5b warmed to room temperature added for 30 seconds. Excess buffer was again manually blotted away, followed by another application of 3μl Kif5b. Grids were then placed in an EM GP automated vitrification device (Leica Microsystems) operating at room temperature and 80% humidity, incubated for a further 45 seconds, then blotted and vitrified in liquid ethane.

### Cryo-EM data collection

Cryo-EM data was acquired at the London Consortium for Cryo-EM (LonCEM) on a Krios G3i operating at 300 kV, with a Gatan K3 direct electron detector and a Bioquantum Imaging Filter. Low-dose movies were collected automatically with EPU software with a super-resolution mode sampling of 0.54 Å/pixel, in zero-loss imaging mode with a 20 eV energy-selecting slit. A defocus range of 0.7-2.5 μm was used and the total movie dose was ∼52 e^-^/Å^2^ spread over 36 frames, with electron (e-) counting at 16 e-/physical pixel/second.

### Cryo-EM data processing

Near-identical processing pipelines were used datasets of ADP-bound Kif5b^MoNeXXX^ or ADP- or AMPPNP-bound Kif5b^MoNeIAK^ (Supplementary Fig. 2). Movie frames were binned x 2 during alignment using Relion v4’s implementation of Motioncorr2^59^, to generate unweighted and dose-weighted sums at the physical pixel size of 1.08 Å/pixel. Full dose sums were used for CTF determination in CTFFIND v4.1.14^60^, then dose-weighted sums used in particle picking, processing and generation of the final reconstructions.

MTs were boxed manually in Relion v3.0’s helical mode and then 4 x binned segments extracted with a box separation distance of 82 Å (roughly the MT dimer repeat distance). From this stage, MT segments were processed with a modified and extended version of MiRP^45^ implemented within Relion v3.0. Briefly, a supervised 3D classification was run to undecorated 11-16 protofilament MT references, with the modal class designated within each MT assigned to all segments within that MT. The dominant 13 protofilament class was then selected and aligned to a 12 Å low-pass filtered reference of a 13-protofilament MT decorated with ADP-AlF_4_-bound kinesin-1 motor domains, generated from PDBs codes 5syf and 4hna using ChimeraX’s^61^ ‘molmap’ command. Rough *x*-*y* translations and ψ angles found were then smoothed between adjacent segments along each MT using custom scripts. Rough ϕ (Phi) angles were then determined for segments with a second 3D refinement and the median φ angles for each MT were assigned to all segments in a given MT. Following another round of smoothing *xy* translations and Euler angle assignments, a fine C1 3D local refinement was then performed with 4x binned segments. ‘Segment averages’ were then generated, by averaging each segment with its 7 adjacent partners within each MT. Using segment averages, assigned φ angles for each MT were checked by supervised 3D-classification without alignment to low-pass filtered kinesin-only references rotated and translated to represent all possible seam positions and αβ-tubulin registers (i.e 26 references for a 13-protofilament MT, with 13 seam positions and their counterparts translated one monomer along the helical axis). Rough final φ angles were assigned according to the modal 3D class of all segments within each MT. Non-binned segments were then reextracted and a fine C1 3D local refinement performed followed by per-particle CTF refinement and Bayesian polishing, each followed by another fine C1 3D local refinement. MT segments were then subjected to an unsupervised 3D classification without alignment using 4 classes and the high-resolution classes selected and inputted into another round of fine C1 3D local refinement.

Symmetry expansion using 13pf MT symmetry parameters determined in Relion was performed. The symmetry expanded coordinates were then used to extract 1 x binned particles (240 pixels at 1.08 Å/pix) focused on the Kif5b^MoNeIAK^ kinesin-tubulin interface of each asymmetric unit. To carry both CTF-refinement and polishing information from whole MT segments to individual asymmetric unit sub-particles, a signal subtraction job, but without signal subtraction applied, was used for this extraction step. Using a reference generated from this extracted asymmetric unit data, a 3D local refinement with a wide circular mask was then performed on these particles. Following this step, a supervised 3D classification without alignment was performed, using references with or without a MT-associated kinesin head. The kinesin-decorated class was selected and then either processed further (for Kif5b^MoNeXXX^ data and Kif5b^MoNeIAK^ data when combining one and two-headed states for maximum resolution of the MT-bound head) or re-extracted focused on the tail density in each asymmetric unit (for separate processing of Kif5b^MoNeIAK^ one and two-headed states focused on the tail region). Particles focused on the kinesin-tubulin interface were subjected to a final local 3D refinement using a soft mask inclusive of the tubulin dimer and MT-associated motor domain. Kif5b^MoNeIAK^ particles extracted focused on the tail were subjected to an unsupervised 3D classification without alignment using a soft mask inclusive of the distal motor head. Two classes one with and one without the second motor head were found. The one-headed class was then subjected to a final local 3D refinement using a soft mask inclusive of the tubulin dimer and MT-associated motor domain. The two-headed class was then subjected to a final local 3D refinement using a soft mask inclusive of either tubulin and both motor domains or just both motor domains without tubulin.

Global resolutions were estimated at the gold-standard FSC 0.143 cut-off (noise-substitution test-corrected), using soft masks around the refined density. Local resolutions were estimated using Relion’s inbuilt software. Reconstructions used for model building, refinement and display were locally sharpened using DeepEMhancer^62^, Relion’s local resolution filtering or LocScale^63^. Final particle numbers are shown in Table 1.

**Table 1.** Cryo-EM data collection, refinement and validation statistics.

|  | MT-Kif5b<sup>MoNeIAK</sup> -<br>AMPPNP<br>one-headed class<br>(EMD-19174,<br>PDB 8RHB) | MT-Kif5b<sup>MoNeIAK</sup> -<br>-AMPPNP<br>two-headed class<br>(EMD-19176,<br>PDB 8RHH) | MT-Kif5b<sup>MoNeIAK</sup> -<br>-ADP<br>one-headed class<br>(EMD-19188<br>PDB 8RIK) | MT-Kif5b<sup>MoNeIAK</sup> -<br>-ADP<br>two-headed class<br>(EMD-19192<br>PDB 8RIZ) |
| --- | --- | --- | --- | --- |
| **Data collection and processing** |  |  |  |  |
| Magnification (actual) | 46,168 | 46,168 | 46,168 | 46,168 |
| Voltage (kV) | 300kV | 300kV | 300kV | 300kV |
| Electron exposure<br>(e<sup>-</sup>/Å<sup>2</sup>) | 52 | 52 | 52 | 52 |
| Defocus range (μm) | -0.7 to -2.5μm | -0.7 to -2.5μm | -0.7 to -2.5μm | -0.7 to -2.5μm |
| Pixel size (Å) | 1.083 Å | 1.083 Å | 1.083 Å | 1.083 Å |
| Symmetry imposed* | Pseudo-helical | Pseudo-helical | Pseudo-helical | Pseudo-helical |
| Initial MT segment<br>particles<sup>a</sup>, (#) | 116,510 | 116,510 | 40,669 | 40,669 |
| 13pf MT segment<br>particles<sup>a</sup>, (#) | 94,840 | 94,840 | 22,594 | 22,594 |
| Initial single particles<br>used<sup>a</sup>, (#) | 1,204,788 | 1,204,788 | 288,184 | 288,184 |
| Final single particles<br>used<sup>a</sup>, (#) | 633,952 | 570,836 | 110,731 | 39,512 |
| Map resolution (Å)<sup>b</sup> | 3.0 Å | 3.0 Å, 3.4 Å | 3.1 Å | 3.6 Å, 3.9 Å |
| FSC threshold<sup>c</sup> | FSC 0.143 | FSC 0.143 | FSC 0.143 | FSC 0.143 |
| Map resolution range<br>(Å)<sup>b</sup> | 2.9-3.6 Å | 2.9-5.6 Å, 3.5-4.1 Å | 2.9-4.1 Å | 3.3-9.3 Å, 3.6-7.8 Å |
| **Refinement** |  |  |  |  |
| Initial models used | 5SYF, 4HNA,<br>2Y65 | 5SYF, 4HNA,<br>2Y65, 1VFV | 5SYF, 4LNU,<br>2Y65 | 5SYF, 4LNU,<br>2Y65, 1BG2 |
| Refinement resolution<br>(Å) | 3.0 | 3.0 | 3.1 | 3.6 |
| **Model composition** |  |  |  |  |
| Nonhydrogen atoms | 9,495 | 12,195 | 9,332 | 11,828 |
| Protein residues | 1,180 | 1,508 | 1,157 | 1,466 |
| Ligands | 6 | 8 | 5 | 7 |
| **R.m.s. deviations** |  |  |  |  |
| Bond lengths (Å) | 0.003 | 0.003 | 0.003 | 0.002 |
| Bond angles (°) | 0.66 | 0.71 | 0.54 | 0.54 |
| **Validation<sup>d</sup>** |  |  |  |  |
| MolProbity score | 1.58 | 1.63 | 1.48 | 1.53 |
| Clashscore | 7.6 | 8.09 | 4.72 | 7.78 |
| Poor rotamers (%) | 0 | 0.22 | 0 | 0.15 |
| **Ramachandran plot<sup>d</sup>** |  |  |  |  |
| Favored (%) | 97.03 | 96.88 | 96.46 | 97.48 |
| Allowed (%) | 2.97 | 3.12 | 3.54 | 2.52 |
| Disallowed (%) | 0 | 0 | 0 | 0 |

|  | MT-Kif5b<sup>MoNeIAK</sup> -<br>AMPPNP<br>all data | MT-Kif5b<sup>MoNeIAK</sup> -<br>ADP<br>all data | MT-Kif5b<sup>MoNeXXX</sup> -<br>ADP<br>(EMD-51477,<br>PDB 9GNQ) |
| --- | --- | --- | --- |
| **Data collection and processing** |  |  |  |
| Magnification | 46,168 | 46,168 | 46,168 |
| Voltage (kV) | 300Kv | 300Kv | 300Kv |
| Electron exposure (e<sup>-</sup>/Å<sup>2</sup>) | 52 | 52 | 52 |
| Defocus range (μm) | -0.7 to -2.5μm | -0.7 to -2.5μm | -0.7 to -2.5μm |
| Pixel size (Å) | 1.083Å | 1.083Å | 1.083 Å |
| Symmetry imposed* | Pseudo-helical | Pseudo-helical | Pseudo-helical |
| Initial MT segment particles<sup>a</sup>, (#) | 116,510 | 40,669 | 125,234 |
| 13pf MT segment particles<sup>a</sup> (#) | 94,840 | 22,594 | 89,119 |
| Initial single particles used<sup>a</sup> (#) | 1,204,788 | 288,184 | 384,502 |
| Final single particles used<sup>a</sup> (#) | 1,204,788 | 150,243 | 268,199 |
| Map resolution (Å)<sup>b</sup> | 2.9 Å | 3.0 Å | 2.9 Å |
| FSC threshold<sup>c</sup> | FSC 0.143 | FSC 0.143 | FSC 0.143 |
| Map resolution range (Å)<sup>b</sup> | 2.8-3.4 Å | 2.9-4.1 Å | 2.8-3.6 Å |
| **Refinement** |  |  |  |
| Initial models used | n/a | n/a | 5SYF, 4LNU, 2Y65 |
| Refinement resolution (Å) | n/a | n/a | 2.9 Å |
| Model composition |  |  |  |
| Nonhydrogen atoms | n/a | n/a | 9,309 |
| Protein residues | n/a | n/a | 1,156 |
| Ligands | n/a | n/a | 5 |
| R.m.s. deviations |  |  |  |
| Bond lengths (Å) | n/a | n/a | 0.005 |
| Bond angles (°) | n/a | n/a | 0.58 |
| Validation<sup>d</sup> |  |  |  |
| MolProbity score | n/a | n/a | 1.42 |
| Clashscore | n/a | n/a | 5.07 |
| Poor rotamers (%) | n/a | n/a | 0.99 |
| Ramachandran plot<sup>d</sup> |  |  |  |
| Favored (%) | n/a | n/a | 97.15 |
| Allowed (%) | n/a | n/a | 2.76 |
| Disallowed (%) | n/a | n/a | 0.09 |

<sup>a</sup>MTs exhibit pseudo-helical symmetry due to a symmetry break at the seam. 13pf pseudo-helical MT segments were selected and processed, then single particles for each unique asymmetric unit produced using symmetry expansion (see methods).
<sup>b</sup>Two-headed classes were refined using either a mask inclusive of both kinesin heads and tubulin (1<sup>st</sup> stated range), or both kinesin heads alone (2<sup>nd</sup> stated range).
<sup>c</sup>The FSC curves for all reconstructions were subjected to the gold-standard noise-substitution test for overfitting<sup>1</sup>, implemented in Relion. The resolution value at the noise-substitution test-corrected FSC 0.143 criterion is shown.
<sup>d</sup>As defined by the MolProbity<sup>2</sup> validation server.

### Pseudo-atomic model building

Initial homology models of AMPPNP or ADP-bound human Kif5b^MoNeIAK^ and Kif5b^MoNeXXX^ MT associated heads were created using MODELLER^64^ and template X-ray structures of a kinesin-1 motor domain bound to AMPPNP (PDB code 4hna) or ADP (PDB code 4lnu), respectively. A taxol-bound tubulin dimer (PDB code 5syf) was added to these models. To make the Kif5b^MoNeIAK^ two-headed state with ADP, a 2^nd^ head was created using MODELLER and a combination of ADP-bound kinesin-1 structures (PDB codes 1bg2 and 2y65). To make the Kif5b^MoNeIAK^ two-headed state with AMPPNP, a 2^nd^ head was created using MODELLER and a combination of AMPPNP-bound kinesin-3 (PDB code 1vfv) and ADP-bound kinesin-1 (PDB code 2y65) motor domain crystal structures. The tail was modelled based on the *Drosophila melanogaster* kinesin-1 autoinhibited dimer crystal structure (PDB code 2y65). After trimming the models to remove disordered regions not represented by EM density, these models were rigid-fitted into cryo-EM density with the ‘fit-in-map’ ChimeraX tool. Iterative rounds of Coot^65^ and real space refinement in Phenix^66^ were then performed. For Kif5b^MoNeIAK^ single-headed states the 2^nd^ head was removed, and part of the tail deleted, followed by a further round of real space refinement. Cryo-EM density and model analysis and display used ChimeraX. Model refinement statistics are presented in (Supplementary Table 1).

## Data availability

MT-bound Kif5b^MoNeXXX^ and Kif5b^MoNeIAK^ one and two-headed asymmetric unit reconstructions we deposited within the Electron Microscopy Data Bank (EMDB) alongside corresponding atomic models deposited in the Protein Data Bank (PDB). EMDB and PDB accession codes are shared in (Supplementary Table 1).

## Acknowledgements

J.A. and T.F. are supported by the Biotechnology and Biological Sciences Research Council (BBSRC) grant BB/V006568/1. M.S.C. was supported by the BBSRC grant BB/S000828/1 awarded to R.A.S. E.P. and L.S.P. are supported by the Italian Ministry for Universities and Research (MUR) grants PRIN-PNRR (P2022LSH5A) and PRIN (2022ERB7SL), respectively, awarded to R.A.S. We acknowledge the contribution of two Master students in the R.A.S. group, Miss Miral Tariq and Mr Henry Cornish, in the early phase of this work. The London Consortium for Electron Microscopy (LonCEM) is supported by the Wellcome Trust grant 206175/Z/17/Z and its partner institutes.

## Contributions

J.A., M.S.C., T.F., E.P., S.L.P. carried out the experiments. J.A. and R.A.S. conceived the study, designed the experiments, interpreted the results, and wrote the paper.

## Ethics declaration

Competing interests: the authors declare no competing interests.

## Supplementary Information

**Supplementary Fig. 1.**
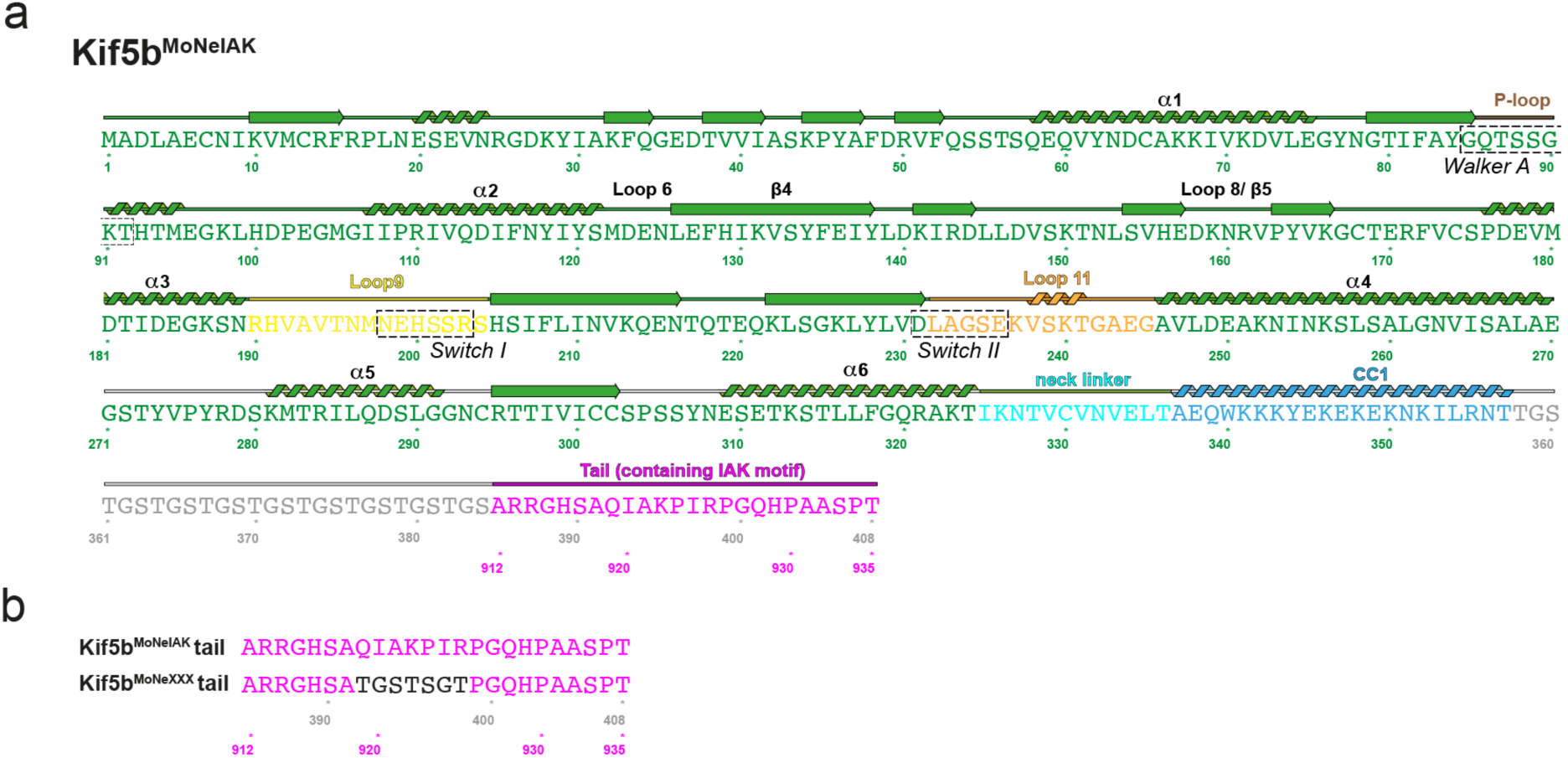
Kif5b^MoNeIAK^ and Kif5b^MoNeXXX^ chimeras and color scheme. **a**. Schematic of the Kif5b^MoNeIAK^ construct showing the motor domain’s sequence, secondary structure elements (SSE), motifs, and coloring scheme used in figures. SSE annotation is based on the structure of our MT-bound structure in the presence of AMPPNP. **b**. Sequence alignment of the Kif5b^MoNeIAK^ and Kif5b^MoNeXXX^ tail regions, (sequence numbering as in panel **a**), with Kif5b^MoNeXXX^ IAK-motif substitutions shown in black.

**Supplementary Fig. 2.**
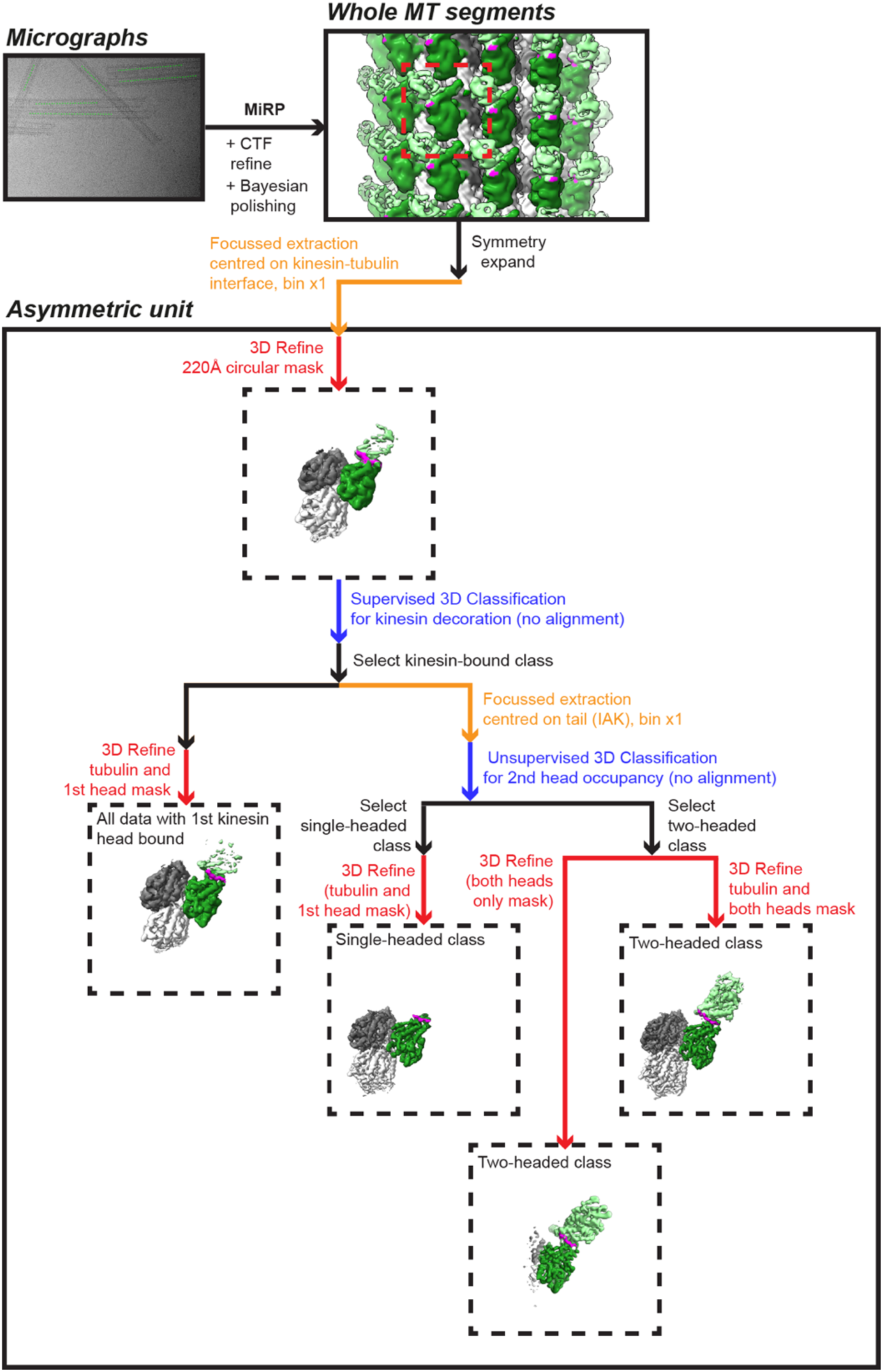
Schematic showing the processing strategy for MT-associated Kif5b^MoNeIAK^ and Kif5b^MoNeXXX^. A full explanation is given in the Methods section, but in brief, micrographs were picked manually in Relion and then whole MT segments processed with MiRP^3^ running in Relion v3.0. Symmetry expansion and focused extraction then allowed us to perform focused refinements and classifications on different portions of the asymmetric unit in using smaller, less memory-intensive box sizes. Focused refinement steps are shown in red, focused extraction steps in shown in orange, focused classification steps shown in blue and other processing steps shown in black.

**Supplementary Fig. 3.**
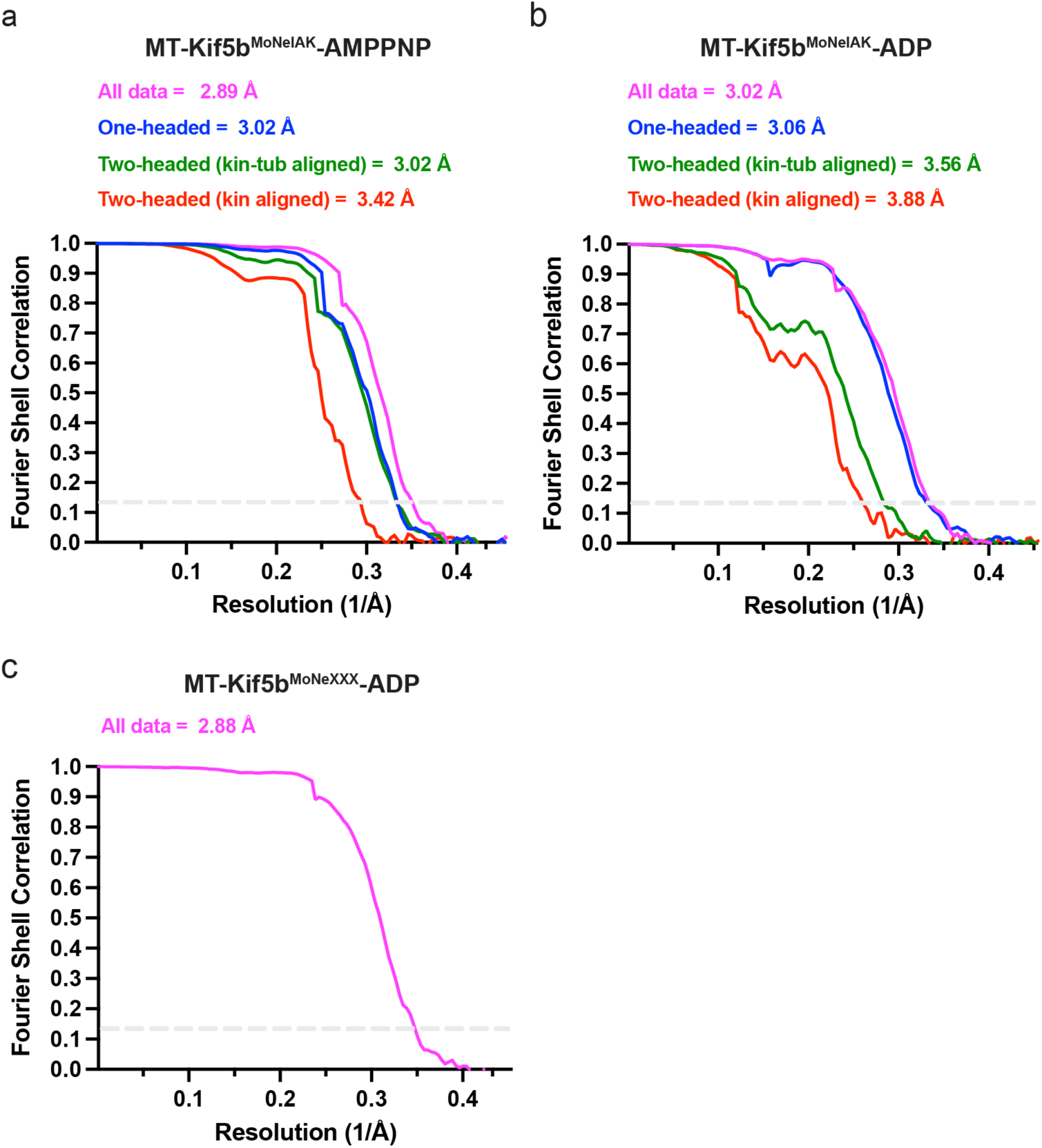
Global FSC curves for all reconstructions. **a,b,c.** Noise-substitution test-corrected FSC curves (derived from Relion v3.0) are shown for focused asymmetric unit reconstructions from (**a**) MT-Kif5b^MoNeIAK^-AMPPNP, (**b**) MT-Kif5b^MoNeIAK^-ADP and (**c**) MT-Kif5b^MoNeXXX^-ADP datasets. For all datasets, reconstructions were either of: *i*) all data refined using a tubulin plus kinesin motor domain-inclusive mask (magenta); *ii*) the one-headed state refined using a tubulin+kinesin motor domain-inclusive mask (blue); the two-headed state refined using either *iii*) a tubulin+both kinesin motor domains-inclusive mask (green) or *iv*) both kinesin motor domains-inclusive mask (excluding tubulin, red). Soft loose masks were used in focused refinements and the same masks used in FSC calculations. Global resolutions in Å are reported at the gold-standard 0.143 FSC cut-off criterion for the noise-substitution test-corrected FSC curves.

**Supplementary Fig. 4.**
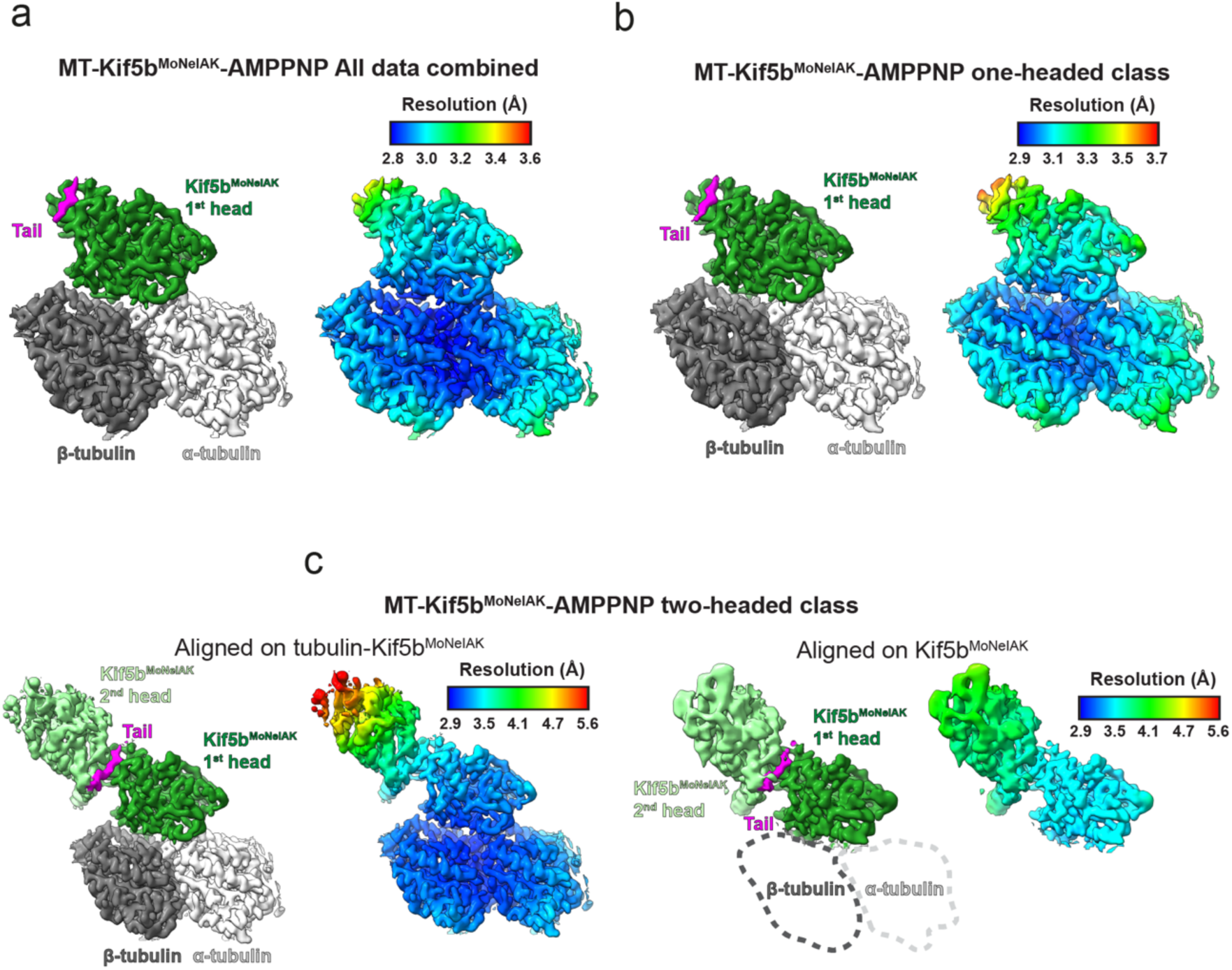
Local resolution estimates for MT-Kif5b^MoNeIAK^-AMPPNP reconstructions. **a,b,c.** Density for asymmetric units is colored either according to segmentation into kinesin motor domain, tubulin and tail densities (left-hand panels) or local resolution (right-hand panels). Similar views are shown for (**a**) all data refined using a tubulin plus kinesin motor domain-inclusive mask, (**b**) the one-headed state refined using a tubulin+kinesin motor domain-inclusive mask and (**c**) the two-headed state refined using either a tubulin+both kinesin motor domains-inclusive mask (left hand panels) or both kinesin motor domains-inclusive mask excluding tubulin whose location is indicated by dashed shapes (right hand panels). Unfiltered/unsharpened final reconstructions are shown with density zoned according to focused masks used for refinement. Local resolutions were estimated with Relion v3.0’s inbuilt software. Color keys are provided above the local-resolution density depictions.

**Supplementary Fig. 5.**
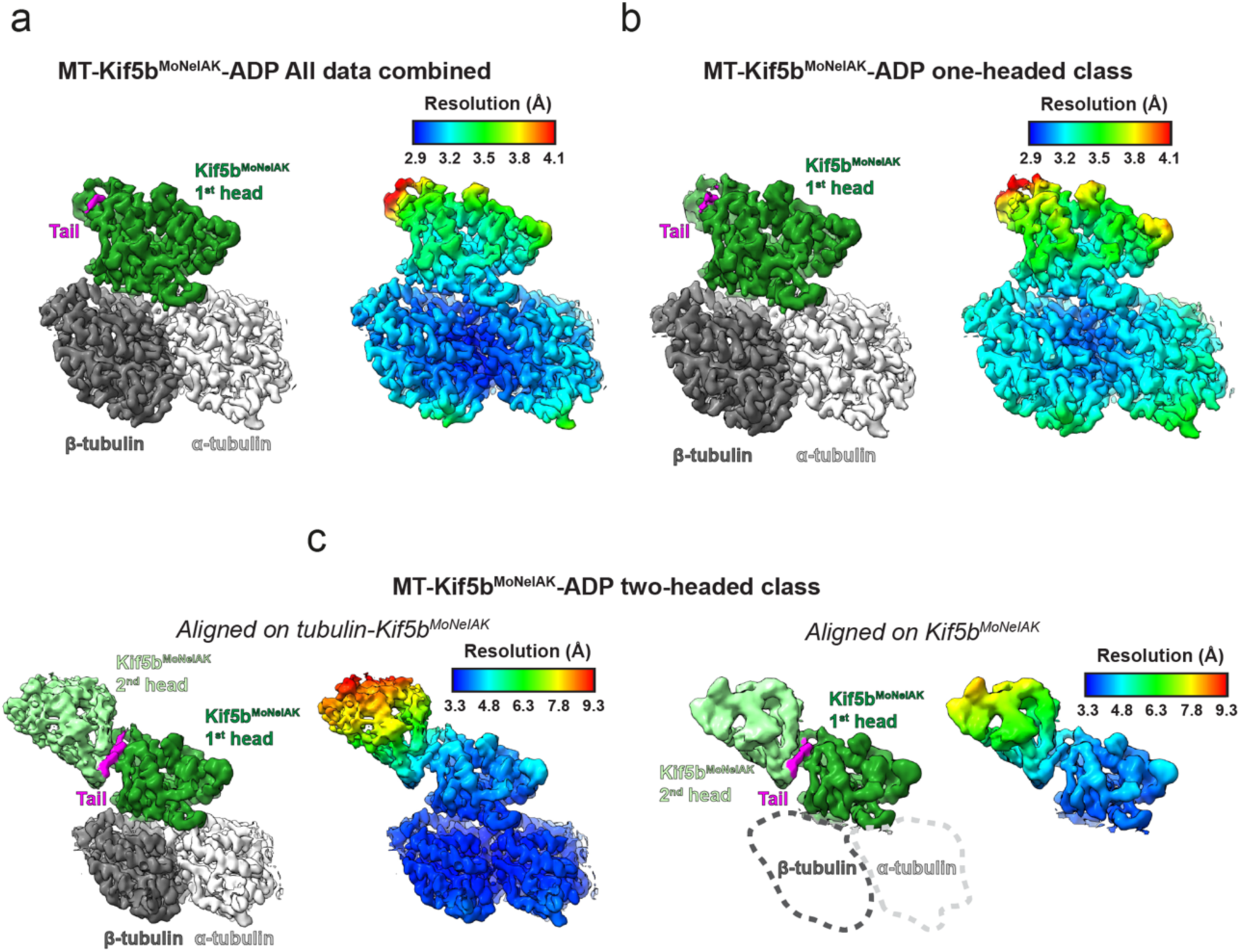
Local resolution estimates for MT-Kif5b^MoNeIAK^-ADP reconstructions. **a,b,c.** Density for asymmetric units is colored either according to segmentation into kinesin motor domain, tubulin and tail densities (left-hand panels) or local resolution (right-hand panels). Similar views are shown for (**a**) all data refined using a tubulin plus kinesin motor domain-inclusive mask, (**b**) the one-headed state refined using a tubulin+kinesin motor domain-inclusive mask and (**c**) the two-headed state refined using either a tubulin+both kinesin motor domains-inclusive mask (left hand panels) or both kinesin motor domains-inclusive mask excluding tubulin which is indicated by dashed shapes (right hand panels). Unfiltered/unsharpened final reconstructions are shown with density zoned according to focused masks used for refinement. Local resolutions were estimated with Relion v3.0’s inbuilt software. Color keys are provided above the local-resolution density depictions.

**Supplementary Fig. 6.**
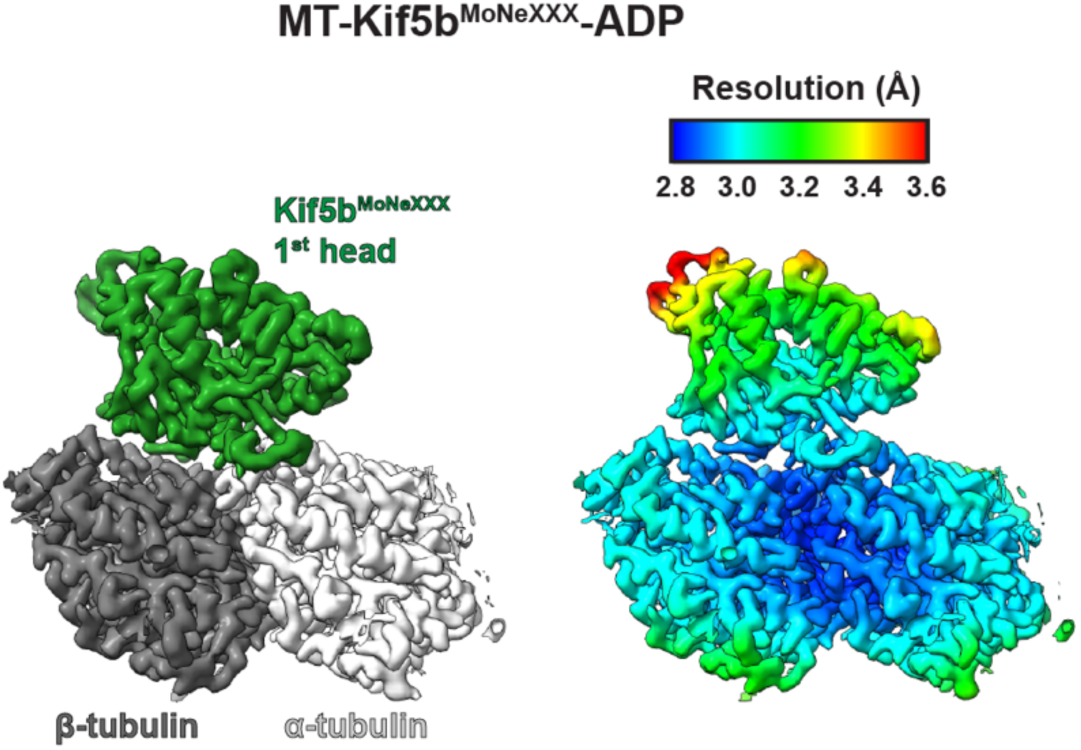
Local resolution estimates for the MT-Kif5b^MoNeXXX^-ADP reconstruction. Density for the asymmetric unit is colored either according to segmentation into kinesin motor domain, tubulin and tail densities (left-hand panel) or local resolution (right-hand panel). The unfiltered/unsharpened final reconstruction is shown with density zoned according to focused masks used for refinement. Local resolution was estimated with Relion v3.0’s inbuilt software. A color key is provided above the local-resolution density depictions.

**Supplementary Fig. 7.**
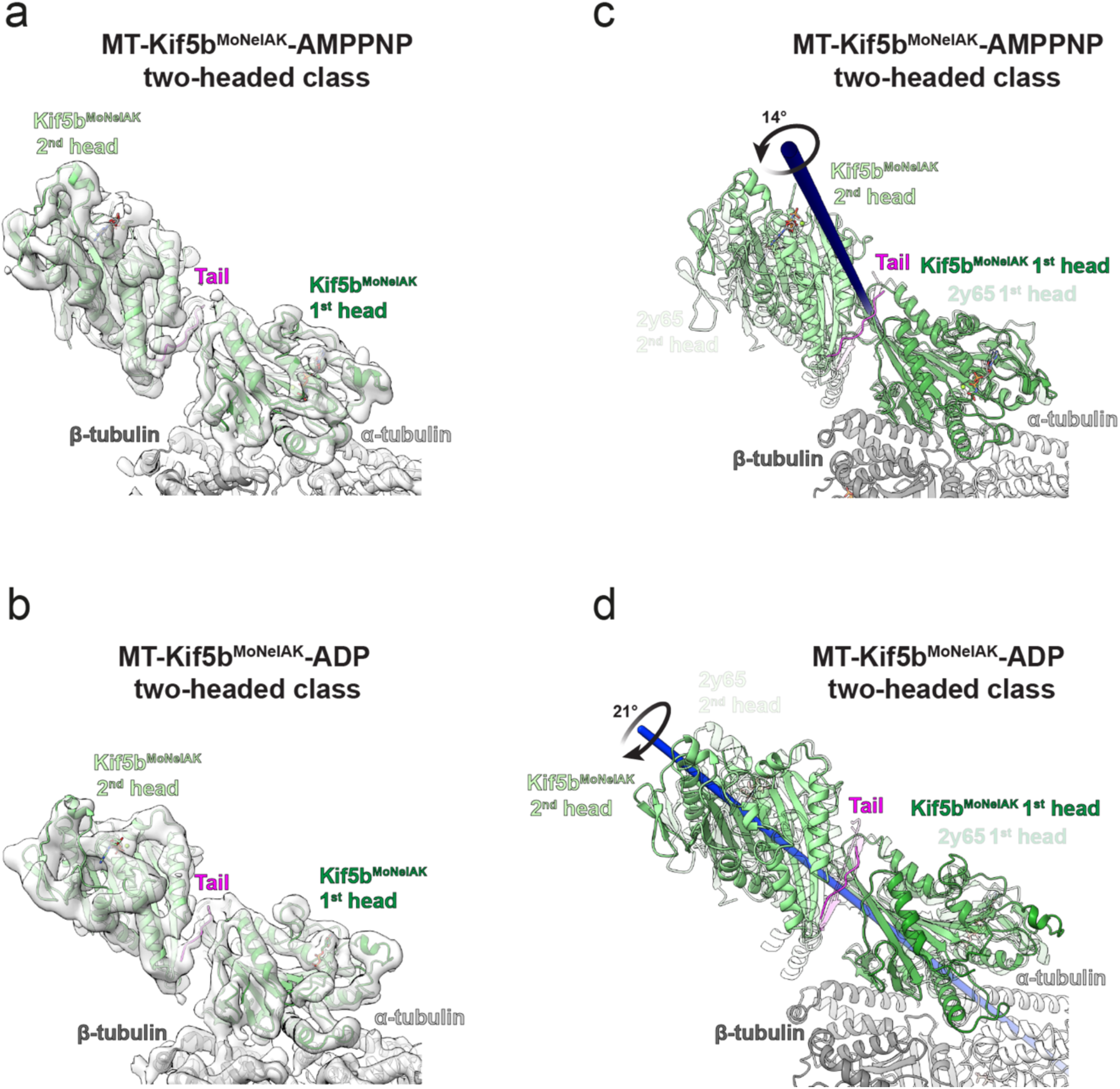
The orientation of the 2^nd^ head relative to the 1^st^ is altered compared to the crystal structure of *Drosophila* kinesin-1 dimer + tail. **a**,**b.** MT-Kif5b^MoNeIAK^ two-headed models bound to (**a**) AMPPNP or (**b**) ADP are shown in their respective cryo-EM densities (transparent grey) low-pass filtered to 6 Å (appropriate for visualization of the whole complex). **c**,**d.** Superimpositions of the X-ray crystallographic structure of *Drosophila melanogaster* kinesin-1 in complex with a tail peptide (semi-transparent, PDB code 2y65)^4^ onto loop 8 and β4 (tail-binding regions) of the MT-associated head of the Kif5b^MoNeIAK^ two-headed models (opaque) bound to either (**c**) AMPPNP (top panel) or (**d**) ADP. The rotation axes (blue) and angles of the 2^nd^ Kif5b^MoNeIAK^ head relative to the 2^nd^ head of the crystallographic model are shown.

**Supplementary Fig. 8.**
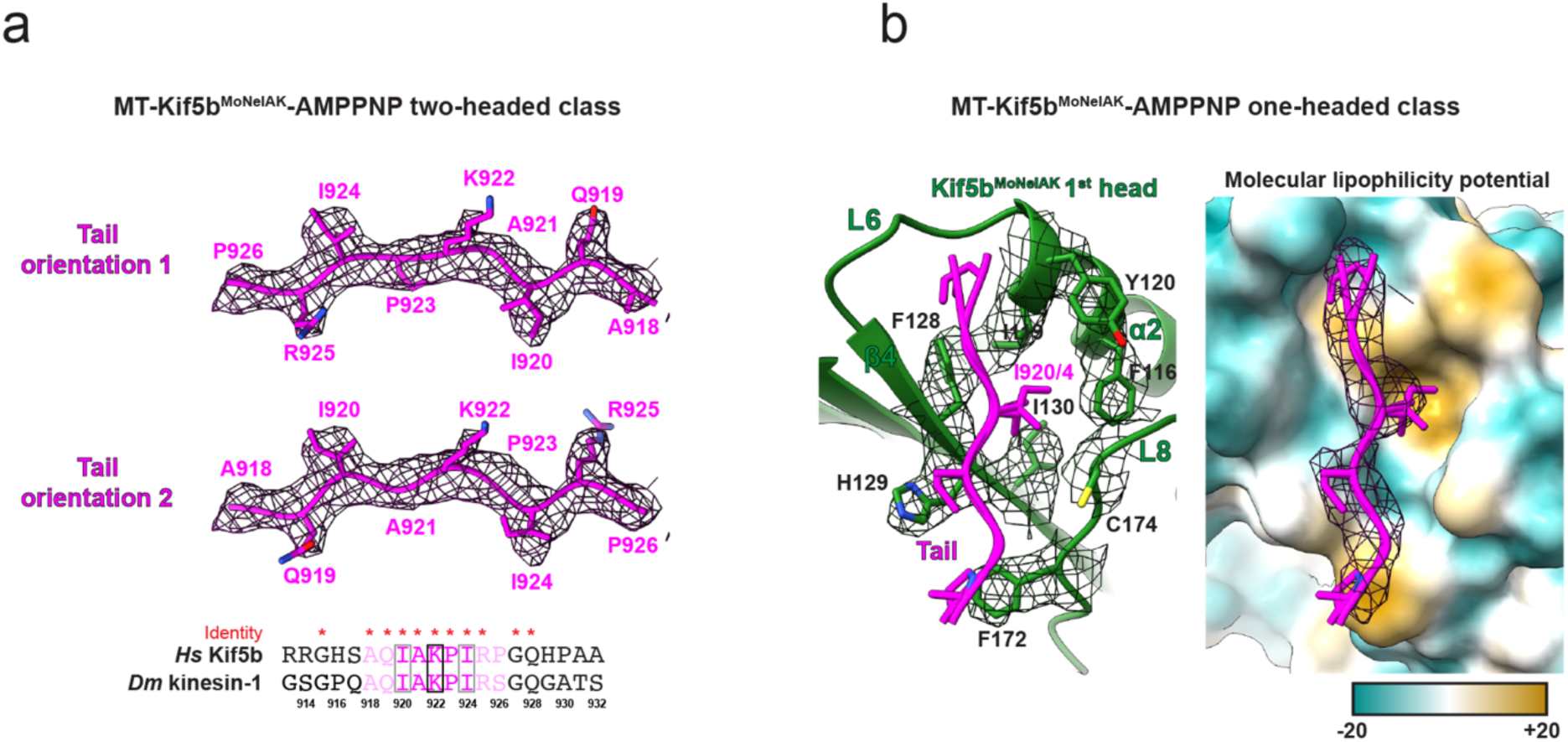
Binding site and mixed polarity of the tail IAK-motif. **a.** Tail density for the two-headed class of MT-Kif5b^MoNeIAK^ in the presence of AMPPNP with two modelled orientations of the pseudo-palindromic tail sequence with amino acid numberings. A sequence alignment of the IAK-motif containing tail region of human Kif5b (*Hs*Kif5B) and *Drosophila melanogaster* kinesin-1 (*Dm*kinesin-1) is shown below, with identical residues illustrated with red asterisks. Light and solid magenta sequence coloring shows residues resolved in the presence of AMPPNP, with residues only resolved in the presence of ADP shown in solid magenta. The central lysine K922 of the pseudo-palindrome is boxed in black, whereas key isoleucine residues I920 and I924 either side are boxed in grey. **b.** The motor domain head (green) binding site of the tail (magenta) is shown on the AMPPNP-bound one-headed state, with focus on the embedding of I920/4 a hydrophobic pocket. The left-hand panel shows the motor domain model binding site, with side chains and density (mesh) for key Kif5b hydrophobic pocket residues associated with the tail. The right-hand panel shows the same view, but with tail density shown in mesh and the motor domain displayed as a surface colored by hydrophobicity (molecular lipophilicity potential calculated by ChimeraX^5^, range −20 to +20).

**Supplementary Fig. 9.**
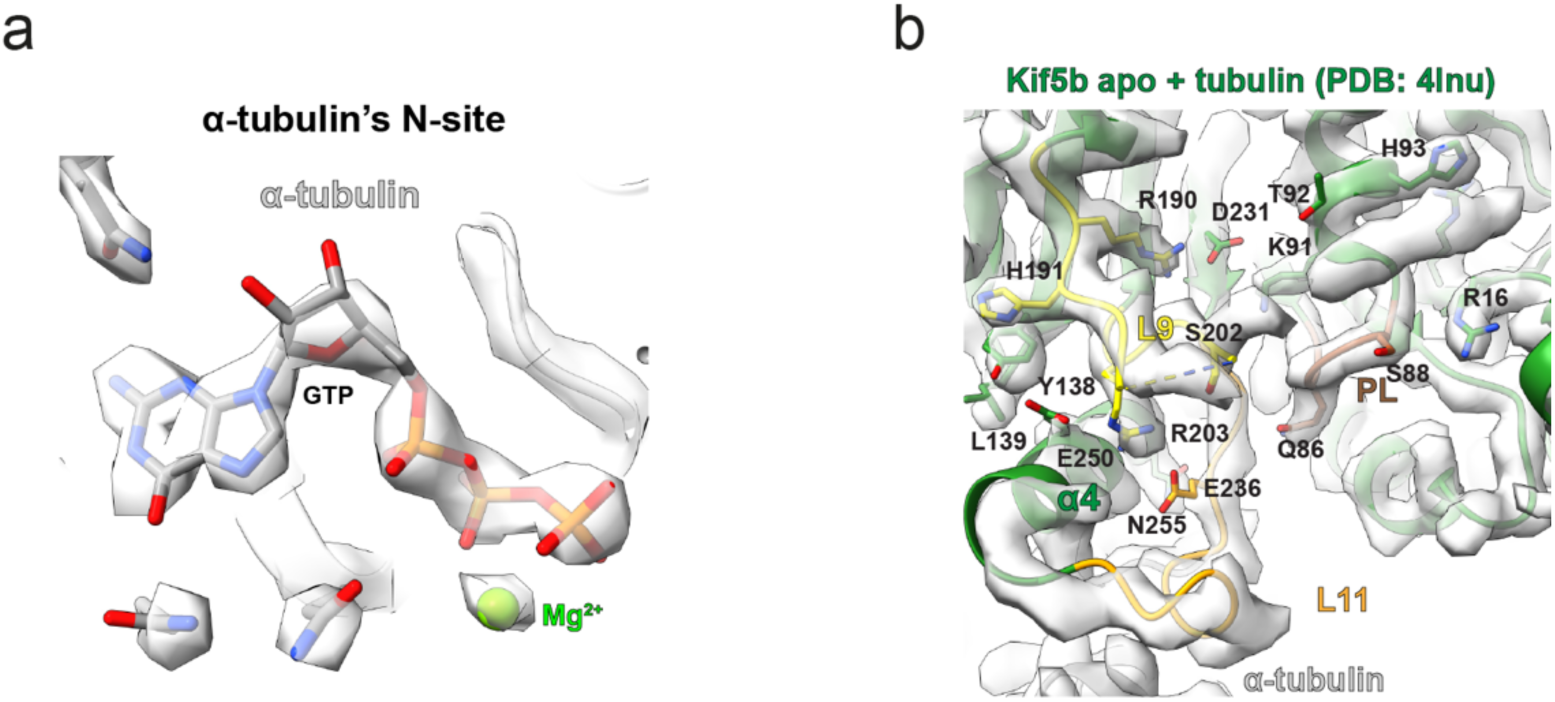
The MT- and ADP-bound Kif5b^MoNeXXX^ complex is apo-like and retains Mg^2+^ in α-tubulin’s N-site. **a.** α-tubulin’s non-exchangeable GTP binding site in the reconstruction from all MT-associated Kif5b^MoNeXXX^-ADP data, showing clear cryo-EM density (semi-transparent grey) for a Mg^2+^ ion associated with GTP. **b.** Overview of the nucleotide pocket and switch-motif containing loops L9 and L11 in the nucleotide-free (apo) structure of Kif5b motor domain on tubulin (PDB code 4lnu)^6^, with side chains for key conserved residues shown. The model is fitted into the cryo-EM density for MT-associated Kif5b^MoNeXXX^-ADP (semi-transparent), showing the high structural similarity (apart from the missing ADP).

**Supplementary Fig. 10.**
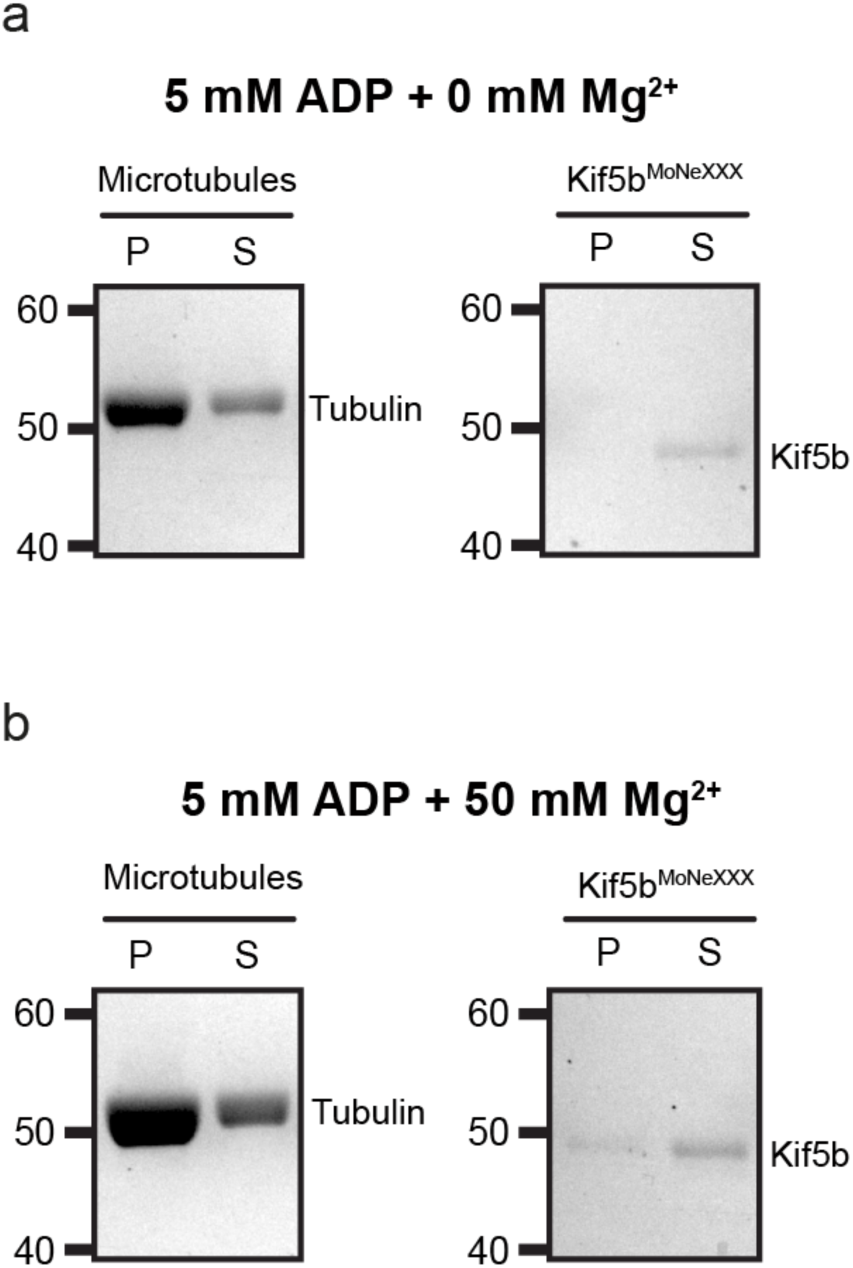
Controls for the co-sedimentation assay shown in Fig. 3b,c of the main text. **a**,**b.** Controls of MT-only or Kif5b^MoNeXXX^ only samples sedimentation with either (**a**) no MgCl_2_ or (**b**) 50 mM MgCl_2_. The experiment was performed in BRB80 buffer with 5 mM ADP and the indicated concentrations of MgCl_2_, with taxol-stabilized MTs at 2 μM (tubulin dimer) and 0.5 μM Kif5b^MoNeXXX^. For conditions without Mg^2+^ we used 10 mM EDTA to remove any possible free ion present. P = Pellet, S = Supernatant.

**Supplementary Fig. 11.**
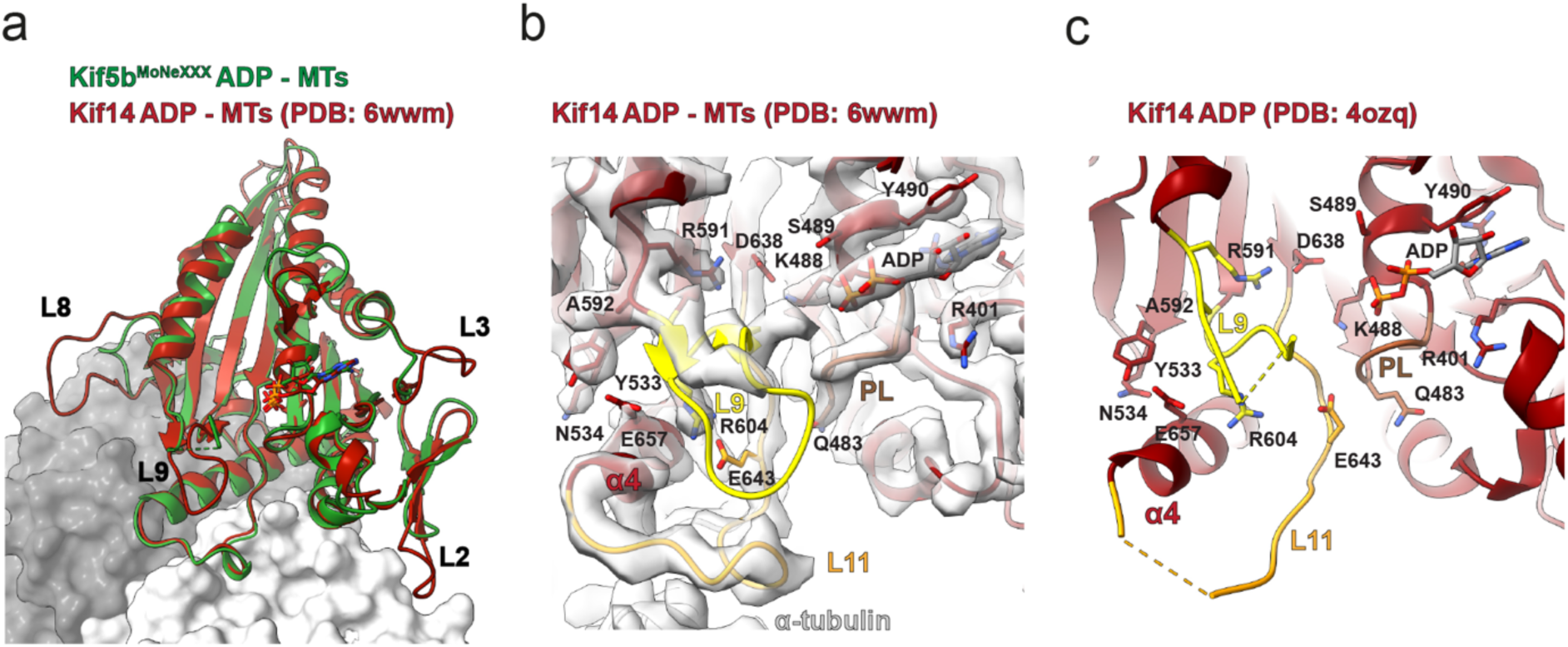
Comparison of Kif14 apo-like ADP states with Kif5b^MoNeXXX^-ADP structure. **a.** ADP-bound motor domains of the Kif5b^MoNeXXX^ (green) and Kif14 (maroon, PDB code 6wwm)^7^ MT-associated models are superimposed, with divergent loops labelled. The tubulin dimer from the Kif5b^MoNeXXX^ asymmetric unit is shown as a grey surface representation. **b**. Overview of the kinesin nucleotide-pocket and switch-motif containing loops L9 and L11, where the Kif14-ADP MT-associated model is fitted into our microtubule-associated Kif5b^MoNeXXX^-ADP cryo-EM density (semi-transparent), showing high similarity apart from L9’s apex and certain divergent amino acid side chains. **c.** A similar view to panel (**b**), but instead showing the structure of Kif14-ADP in the absence of MTs (PDB code 4ozq)^8^ without cryo-EM density. Sequence numberings for Kif14 are shown.

**Supplementary Fig. 12.**
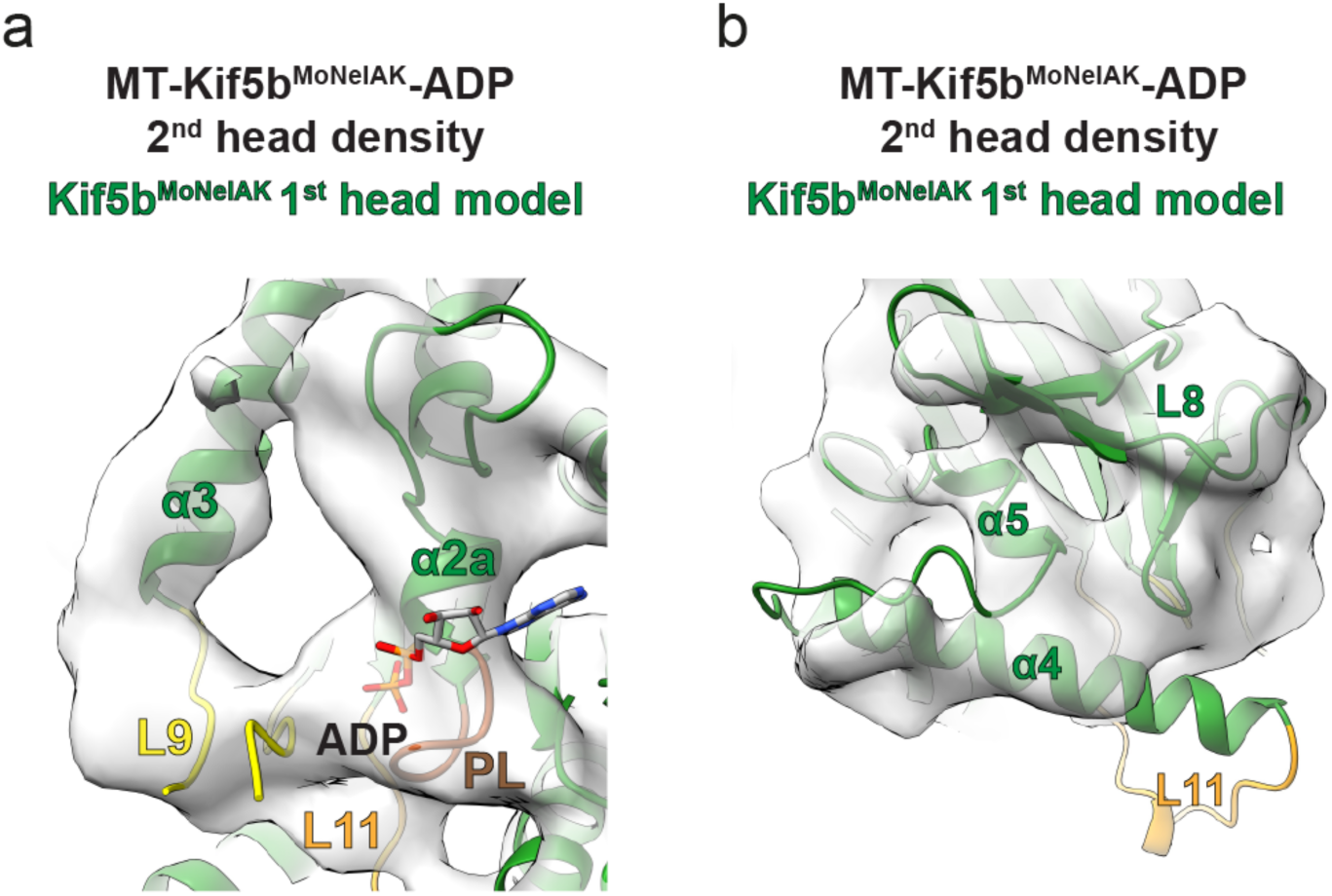
In the presence of ADP, the 2^nd^ head of the Kif5b^MoNeIAK^ two-headed state adopts a distinct conformation compared to the 1^st^ head. Cryo-EM density (semi-transparent, low-pass filtered to 6 Å) is shown for the 2^nd^ (not MT-associated) Kif5b^MoNeIAK^ head in the presence of ADP with views of (**a**) the nucleotide pocket and (**b**), the MT binding elements (including helix-α4 and L11) and the neck linker. The model for the Kif5b^MoNeIAK^ microtubule-associated 1^st^ head is fitted into the density for the 2^nd^ head, showing the two heads adopt distinct conformations (compare with Fig. 4f,g).

**Supplementary Fig. 13.**
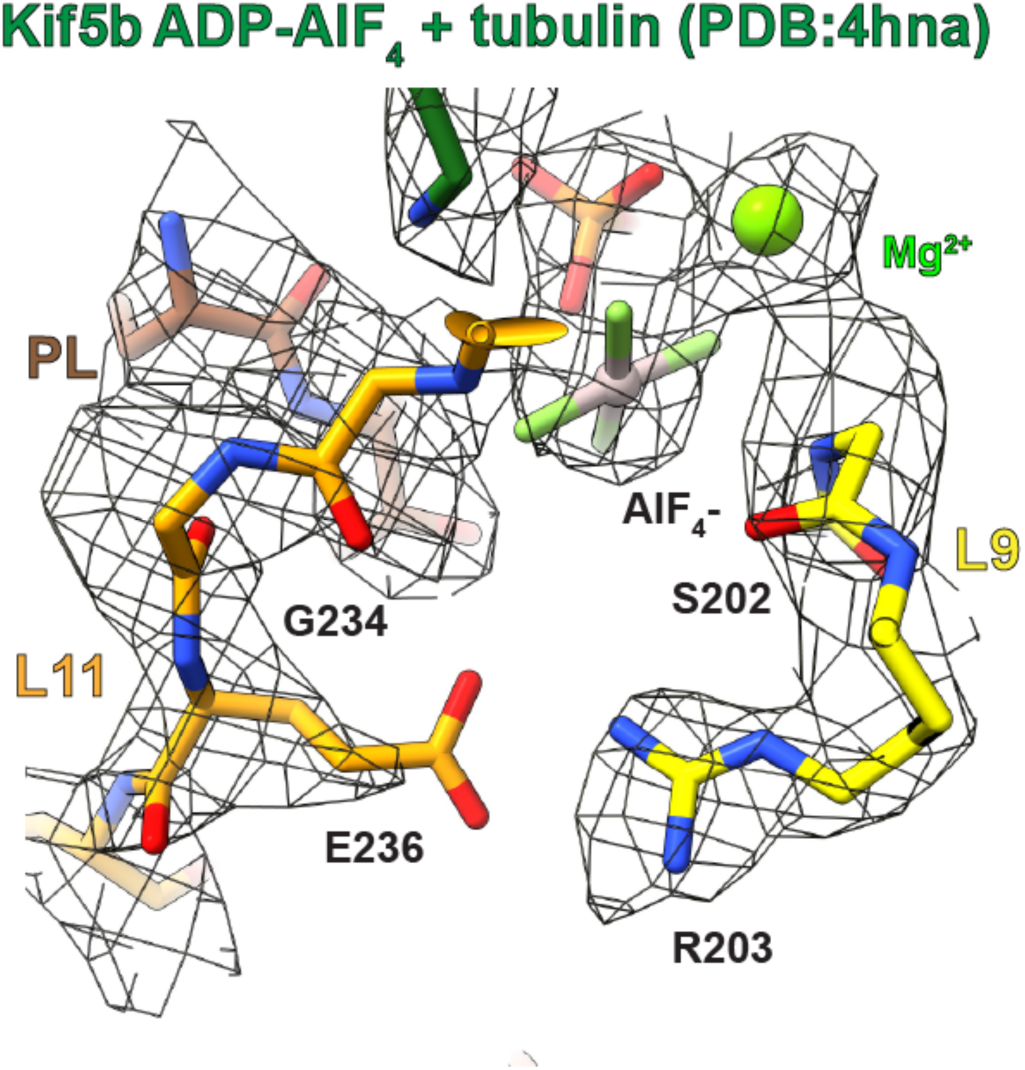
The ATP-like state of nucleotide pocket is indistinguishable in tail-bound and tailless Kif5b. Close up view of residues involved in ATP hydrolysis, with the tubulin and ADP-AlF_4_-bound tailless crystal structure of the Kif5b motor domain (PDB code 4hna)^9^ fitted into density for the MT-associated motor domain of AMPPNP-bound Kif5b^MoNeIAK^ (mesh).

**Supplementary Fig. 14.**
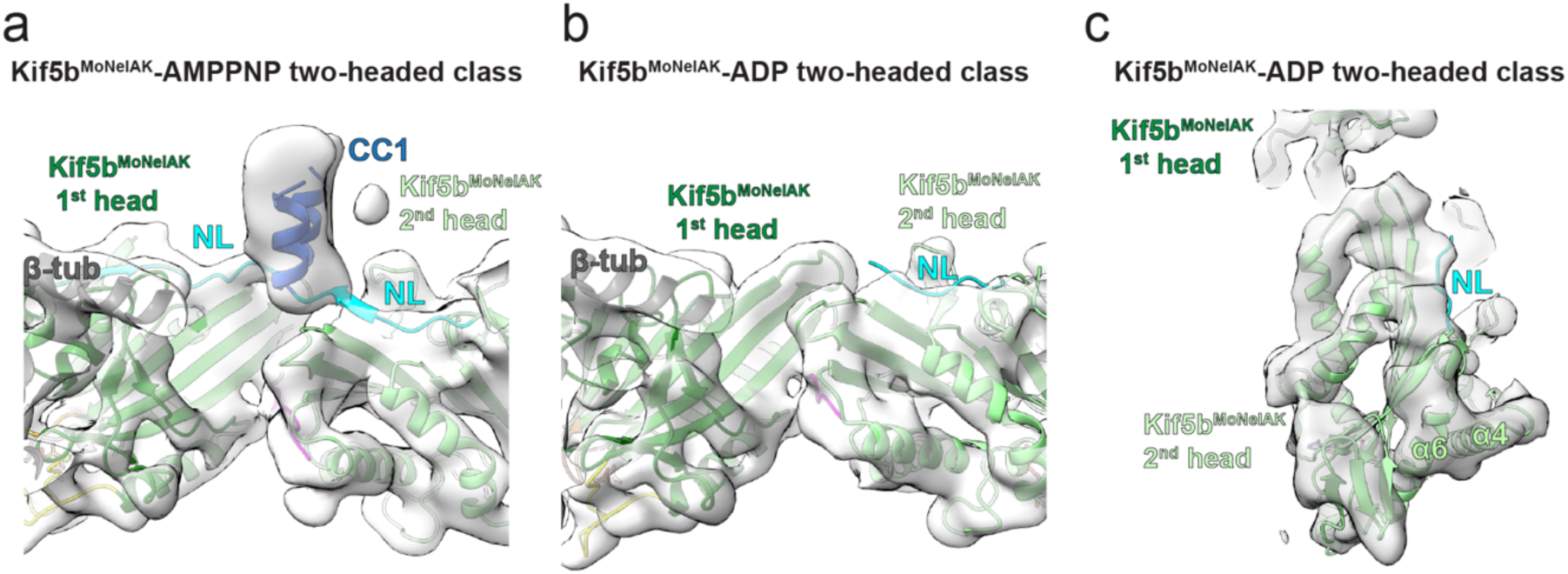
Neck linkers of both heads in the two-headed state only dock in the presence of AMPPNP, such that CC1 is observed. **a**,**b.** Views of the inter-head interface of the Kif5b^MoNeIAK^ two-headed state in the presence of (**a**) AMPPNP or (**b**) ADP, with the corresponding semi-transparent density (low-pass filtered to 6 Å, appropriate for visualization of the whole complex) and fitted models shown. Both neck-linkers (NL) are only docked in the presence of AMPPNP, such that CC1 is stabilized. **c.** Alternative view of the 2^nd^ head in the ADP-bound Kif5b^MoNeIAK^ two-headed state, showing docking of the neck linker.

**Supplementary Fig. 15.**
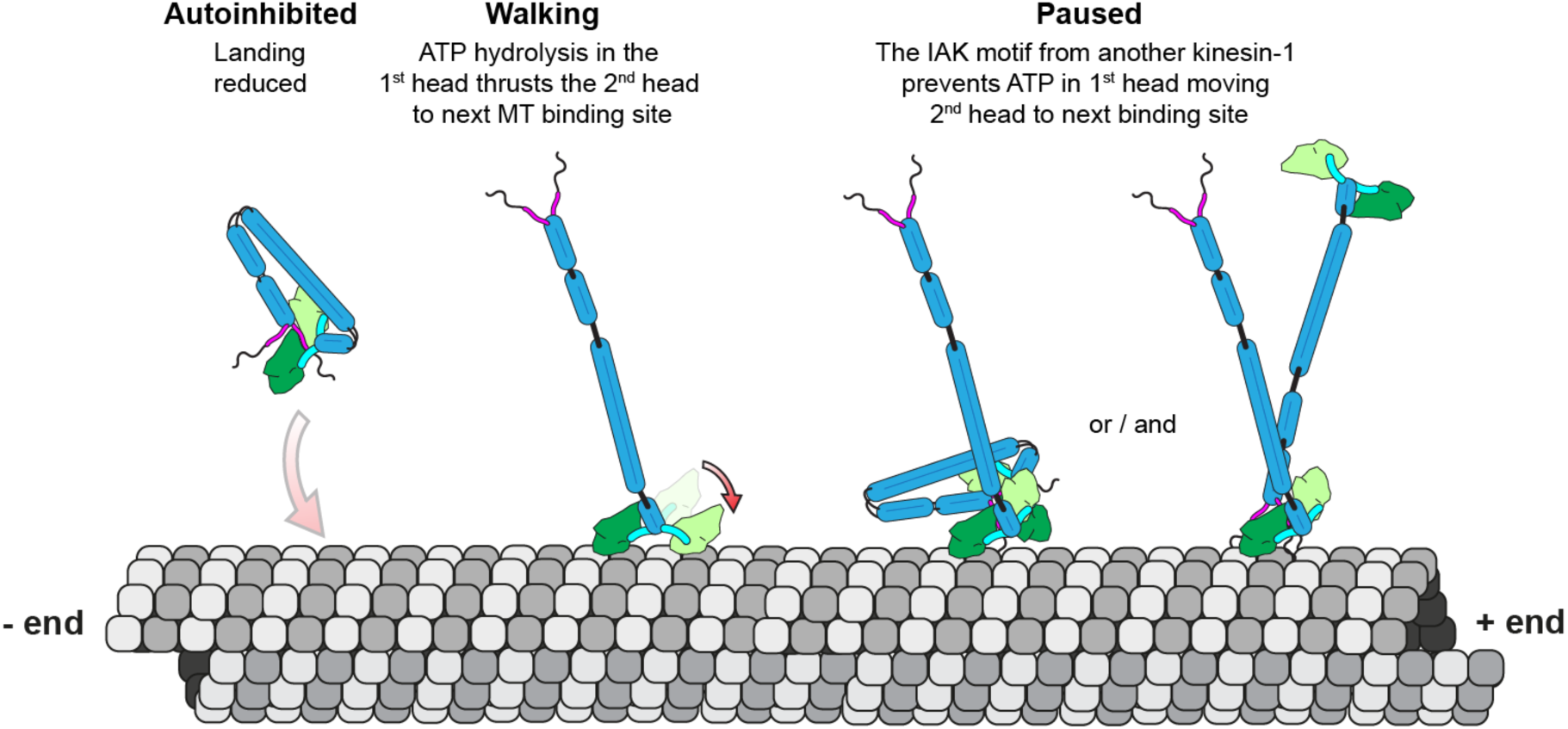
Schematic illustrating mechanisms of kinesin-1 autoinhibition and motor pausing. In the autoinhibited conformation of full-length kinesin-1 landing events are reduced by steric blockage of the MT binding motor domain interface by regions of the coiled coil, with the IAK-containing tail helping to stabilize this conformation. In walking kinesins, ATP binding in the MT-bound head thrusts the second head forward to the next tubulin binding site, however IAK-containing tails from other (either autoinhibited or not) kinesin-1s can lead to motor pausing by locking the second head against the first. The IAK-motif however does not have any effect on the conformation of the MT-bound motor domain.

